# Gray and white matter morphology in substance use disorders: A neuroimaging systematic review and meta-analysis

**DOI:** 10.1101/2020.05.29.122812

**Authors:** Victor Pando-Naude, Sebastian Toxto, Sofia Fernandez-Lozano, E. Christine Parsons, Sarael Alcauter, Eduardo A. Garza-Villarreal

**Affiliations:** Department of Clinical Medicine, Center for Music in the Brain, University of Aarhus, Aarhus, Denmark; Instituto de Neurobiología, Universidad Nacional Autónoma de México (UNAM) campus Juriquilla, Queretaro, Mexico; Instituto Nacional de Psiquiatría “Ramón de la Fuente Muñiz”, Mexico City, Mexico; Facultad de Medicina, Universidad Nacional Autónoma de México (UNAM), Mexico City, Mexico; Facultad de Psicología, Universidad Nacional Autónoma de México (UNAM), Mexico City, Mexico; Department of Clinical Medicine, Interacting Minds Center, University of Aarhus, Aarhus, Denmark; Department of Clinical Medicine, Center of Functionally Integrative Neuroscience, University of Aarhus, Aarhus, Denmark

## Abstract

Substance use disorders (SUDs) are characterized by a compulsion to seek and consume one or more substances of abuse, with a perceived loss of control and negative emotional state. Repeated use of a substance results in synaptic and morphological changes, secondary to toxicity and SUD pathology in the dopamine striato-thalamo-cortical and limbic pathways. These neuroadaptations seem to vary between studies, which could be related to divergent effects of substances, consumption severity or other unknown factors. We therefore identified studies investigating the effects of SUDs using volumetric whole-brain voxel-based morphometry (VBM) in gray (GM) and white matter (WM). We performed a systematic review and meta-analysis of VBM studies using the anatomic likelihood estimation (ALE) method implemented in GingerALE (PROSPERO pre-registration <u>CRD42017071222</u>). Fifty studies met inclusion criteria and were included in the final quantitative meta-analysis, with a total of 538 foci, 88 experiments and 4370 participants. We found convergence and divergence in brain regions and volume effects (higher vs lower volume) in GM and WM depending on the severity of consumption pattern and type of substance. Convergent pathology was evident across substances in GM of the insula, anterior cingulate cortex, putamen, and thalamus, and in WM of the thalamic radiation and internal capsule bundle. Divergent pathology between occasional use (cortical pathology) and addiction (cortical-subcortical pathology) provides evidence of a possible top-down neuroadaptation. Our findings indicate distinctive brain morphometry alterations in SUDs, which may inform our understanding of disease progression and ultimately therapeutic approaches.

## 1. Introduction

Substance use disorders (SUDs) refer to a wide range of alterations produced by the consumption of abuse substances or drugs. According to the Diagnostic and Statistical Manual of Mental Disorders (DSM-V)^1^, these substances include: alcohol, caffeine, cannabis, hallucinogens, inhalants, opioids, sedatives, hypnotics and anxiolytics, stimulants, tobacco and other. About 275 million people worldwide (around 5.6% of the global population aged 15-64 years) used substances at least once during 2016^2^ and SUDs are recognized as a major public health issue. SUDs affect the reward system, involved in the reinforcement of behaviors and memory, and can lead to chronic use and dependency^3^. Initial substance reward is triggered by dopamine neurons in the ventral tegmental area (VTA), which project to the prefrontal cortex, amygdala and nucleus accumbens (NAc)^4,5^, as well as other ascending monoamine fibers such as norepinephrine and other non-dopaminergic systems within frontal regions^6^.

Additionally, dopaminergic neurons in the substantia nigra pars compacta (SNc) project to the dorsal striatum (nigrostriatal pathway), a pathway implicated in the emergence of habits^7^. A reinforcement effect seems to depend on dopaminergic signaling in the NAc, and chronic use may result in neuroadaptations of the dopamine striato-thalamo-cortical (prefrontal cortex, orbitofrontal cortex and the anterior cingulate cortex) and limbic pathways (amygdala and hippocampus)^4,5^, especially in individuals who may be vulnerable due to genetic and/or environmental factors^8^. Other endogenous systems, such as the opioid and cannabinoid systems, may also contribute to the reinforcement effect by modulating hedonic responses or inhibiting negative affective states^9^.

Ultimately, repeated dopaminergic stimulation from substance use may alter multiple neurotransmitter systems, which in consequence may disrupt neuronal excitability and affect neuroplastic mechanisms (i.e. glutamatergic system)^9,10^. Such substance-induced neuroadaptations are similar to synaptic changes associated with learning, including changes in dendritic morphology and ionotropic glutamate receptors (e.g., AMPA/NMDA), which result in long-term potentiation (LTP) and long-term depression (LTD)^11,12^. These neuroadaptations result in pathological changes in brain morphology that seem to be salient enough to be observed macroscopically with MRI, as shown by neuroimaging studies in humans and animal models^13,14^.

Neuroimaging studies using magnetic resonance imaging (MRI) have specifically shown alterations in grey and white matter in SUDs^15,16^. However, the involved regions vary widely and seem to depend on the type of substance, the consumption severity, the age of first use, the total time of usage, and other associated comorbidities. Morphometric studies investigating the effects of SUDs using volumetric measures such as voxel-based morphometry (VBM), have reported both lower and higher volume in cortical and subcortical gray matter (GM)^17,18^ and white matter (WM)^19,20^. For example, alcohol use disorder (AUD) studies have shown lower GM volume of the amygdala, insula, cingulate gyrus, orbitofrontal gyrus and thalamus^15^, while tobacco use disorder (TUD) studies have shown lower GM volume in thalamus, cingulate gyrus, prefrontal cortex and cerebellum^21^. Cocaine use disorder (CUD) studies have shown lower GM volume in thalamus, insula, orbitofrontal cortex, anterior cingulate cortex, superior temporal cortex and cerebellum.^22^ Conversely, other studies of these same substances have shown higher GM volume in putamen and other nuclei of the basal ganglia^23,24^. Similarly, WM studies have shown different substances affecting different areas in distinct manners. For example, studies of AUD, TUD and CUD have shown lower volume of WM in the corticospinal tract, thalamic radiations and the corpus callosum^21,25–27^. Overall, the structural pathology seems to be both convergent and divergent in terms of localization in between studies, suggesting a complex picture.

Given these findings, it is unclear how SUDs affect brain morphology and how to differentiate between distinct changes caused by substance toxicity and substance dependency^28^. Potential reasons for the variability in findings may include: (1) study definitions (substance use disorder vs addiction vs dependency), (2) polysubstance use, (3) the substance user characteristics, such as age or time of substance use, and (4) methodological differences between morphometric studies (i.e. software used). Thus, a meta-analysis of brain imaging studies provides an opportunity to better understand the mechanisms by which SUDs affect brain morphology, of great interest for treatment follow-up as well as potential marker of therapy success. In this systematic review and meta-analysis of VBM studies, we aimed at finding the overall effect of SUDs in GM and WM volume, and to differentiate the possible mechanisms behind such effects by means of subgroup analyses of the type of substance, consumption severity, age and associated comorbidities.

## 2. Methods and Materials

### 2.1. Literature search, screening and extraction

This systematic review and meta-analysis followed procedures from the Cochrane Handbook for Systematic Reviews^29^, and from the Center for Reviews and Dissemination (https://www.york.ac.uk/crd/). The review protocol was pre-registered in PROSPERO (<u>CRD42017071222</u>). This review was carried in accordance with the PRISMA ^30^. We conducted a systematic literature search in PubMed, Scopus and PsycInfo, using both keywords and MeSH terms for articles published to the end of July 2018. No restrictions were placed on study design, but in order to be eligible for inclusion, the studies must have reported whole-brain VBM analyses. Screening and data extraction were performed using the Covidence tool^31^. The main outcome to extract was any change in gray and/or white matter analyzed using VBM, in stereotactic coordinates, comparing a substance user group and a healthy control group (details in **Supplementary information**).

### 2.2. Quality assessment of MRI studies

Criteria for MRI quality reporting was selected from a set of guidelines for the standardized reporting of MRI studies^32–34^. Such guidelines dictate a more consistent and coherent policy for the reporting of MRI methods to ensure that methods can be understood and replicated.

### 2.3. Analysis and meta-analytic technique

Statistically significant foci from between-group contrasts were extracted and recorded for each study. If necessary, coordinates were converted from Talairach coordinates to MNI space using the Lancaster transform (icbm2tal) incorporated in GingerALE (www.brainmap.org/). All metaanalyses were performed using anatomic likelihood estimation (ALE), implemented in GingerALE, in BrainMap^35^. This method extracts the coordinates from the included studies and tests for anatomical consistency and concordance between the studies. The coordinates are weighted according to the size of the sample (number of participants), and these weightings contribute to form estimates of anatomic likelihood estimation for each intracerebral voxel on a standardized map. This approach treats anatomic foci (input) not as single points, but as spatial probability distributions centered at the given coordinates. Therefore, the algorithm tests to what extent the spatial locations of the foci correlate across independently conducted MRI studies investigating the same construct, and assesses them against a null-distribution of random spatial association between experiments^36^. Statistical significance of the ALE scores was determined by a permutation test using cluster-level inference at p < 0.05 (FWE). As we did not impose any minimum cluster size of supra-threshold voxels, small volume clusters should be interpreted with caution.

The primary outcome was morphological brain differences measured by VBM between substance users (SU) and healthy controls (HC), pooling all substances together, to examine comprehensively the structural changes associated with SUD, independently from the directionality of the effect, the type of substance, type of use or age. To test the directionality of the primary outcome, we pooled coordinates reporting higher volume with substance use (HC < SU) and lower volume with substance use (SU < HC). *Pre-registered* subgroup analyses included age of substance users (adolescents vs adults), consumption severity (addiction vs long-term use vs occasional use), type of substance (alcohol vs tobacco vs cannabis vs cocaine vs stimulants vs opioids vs ketamine vs papers that pooled together substances which we termed polysubstance) and associated comorbidities (pure vs dual). Finally, subgroups were tested for similarity (conjunction) and difference (subtraction) in a contrast analysis. All meta-analyses were conducted separately for GM and WM. We use “addiction” as a synonym for SUD that includes dependency, as the latter definition is fairly recent^1^. Additionally, addiction, long-term use and occasional use could also be regarded as severe-, moderate-, and mild-SUD, respectively.

The meta-analytic results (ALE maps) were visualized using Mango (www.rii.uthscsa.edu/mango) on the MNI152 1mm standard brain, and resulting coordinates were cross-referenced to the Harvard-Oxford Cortical and Subcortical Atlas and the Juelich Histological Atlas via NeuroVault^37^ and FSLeyes^38^, respectively.

Finally, we performed the Fail-Safe N analysis (FSN)^39^ as a measure of robustness against potential publication bias. It refers to the amount of contra-evidence that can be added to a meta-analysis before the results change, and can be obtained for each cluster that survives thresholding in an ALE meta-analysis. A higher FSN indicates more stable results and hence a higher robustness.

## 3. Results

A total of 797 records were identified through database searching, and after removing duplicates, 420 records were initially screened by title and abstract. A total of 140 articles were assessed for eligibility in the full-text screening stage. From these, 50 studies fulfilled criteria for eligibility and were included in both the qualitative and quantitative analyses (**Supplementary Figure 1**).

### 3.1. Characteristics of studies

The characteristics of all studies included in the final meta-analysis are shown in **Table 1**. Fifty studies met inclusion criteria and were included in the final quantitative meta-analysis, with a total of 538 foci and 88 experiments. The total number of participants was 4370, with 49.7% substance users (SU) and 50.3% heathy controls (HC). For the SU subsample, 59% in the addiction group (A), 8% on the long-term use group (LT), and 33% on the occasional use group (O). Alcohol was reported in 20% of studies, tobacco 22%, cocaine 12%, cannabis 12%, opioids 12%, stimulants 6%, ketamine 2%, and polysubstance use 14%. SUD was evaluated by a psychiatrist in 26% of studies, psychologist 8%, clinician 2%, while 64% failed to report the evaluator. The DSM-IV was used in 78% of studies, DSM-V 6%, while 16% failed to report the tool used to diagnose substance use disorder. All of the studies reported change in GM volume (100%), while 15 studies (30%) reported change in WM volume.

**Table 1.**
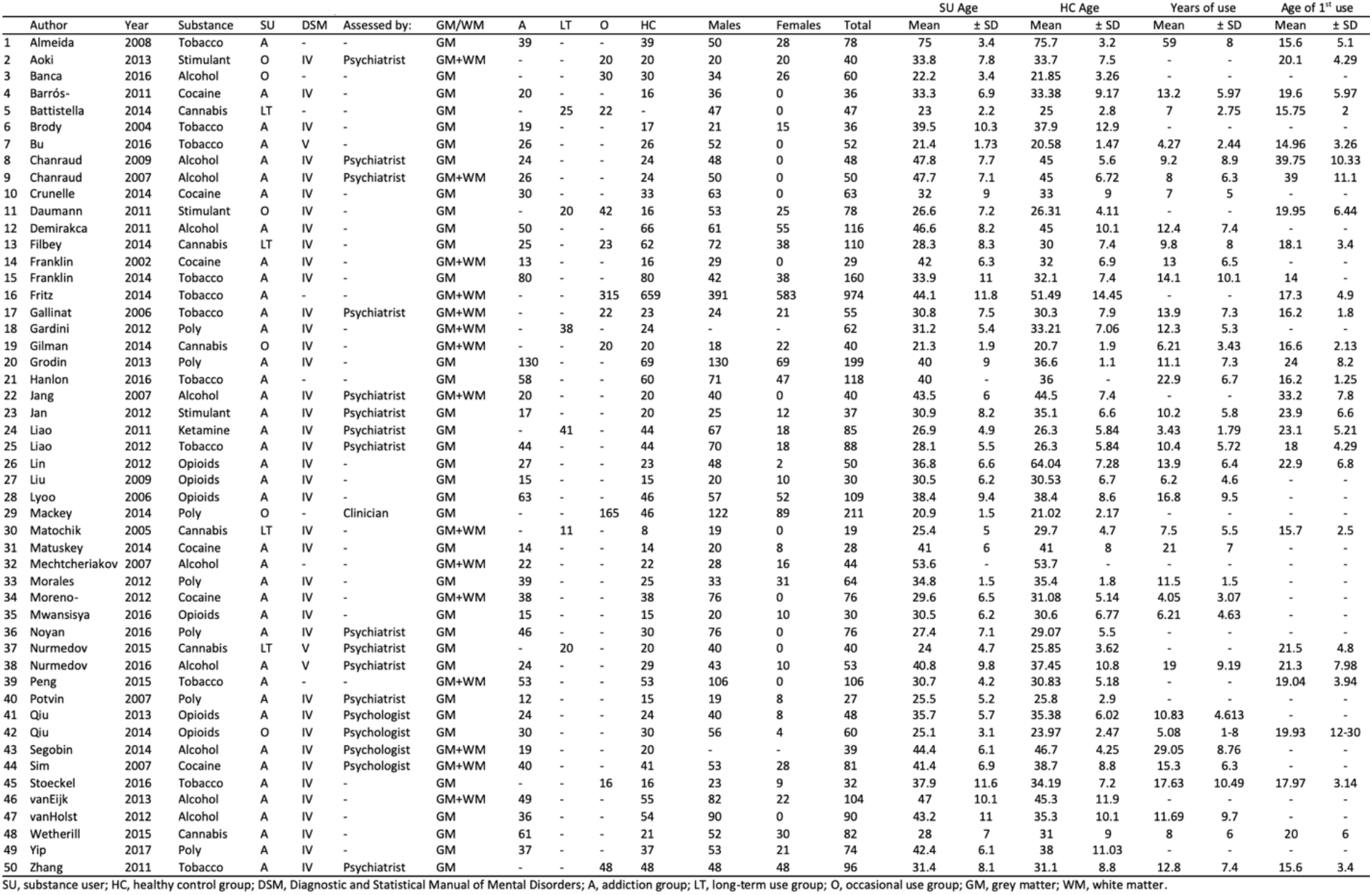
Characteristics of the included in mete-analysis

Neuroimaging data was acquired in either 1.5 T (50%), or 3 T (50%) MRI scanners. Half of the studies were conducted in a Siemens MRI system, others were General Electric (26%), Phillips (22%), and MEDSPEC (2%). Most of the T1w-structural images were acquired using magnetization-prepared rapid acquisition with gradient echo sequence (MPRAGE), in 21 studies (42%), and 1mm^3^-voxel size in 27 studies (54%). VBM analyses were conducted in either SPM^40^ (82%), FSL^38^ (18%) or AFNI^41^ (2%) (**Supplementary Table 1**).

### 3.2. MRI quality

MRI quality of the included studies in the meta-analysis was assessed by a set of guidelines for the standardized reporting of MRI studies^32–34^ (**Supplementary Table 2**). All studies reported the MRI design, software package and image acquisition, processing and analyses. Overall, good MRI practices were performed in the included studies.

### 3.3. Primary outcome

The primary outcome was brain morphological differences measured by VBM between SU and HC, pooling all substances together, and defined as higher or lower volume. First, we included all substances and all reported coordinates and found two clusters in GM: left putamen and left thalamus; and one cluster in WM: right anterior thalamic radiation. Second, the comparison SU < HC (lower volume with use) resulted in 3 GM clusters: left thalamus, left insula and right anterior cingulate cortex; and one WM cluster: right anterior thalamic radiation. Finally, the comparison HC < SU (higher volume with use) resulted in one GM cluster: left putamen; and three WM clusters: right corticospinal tract, left superior longitudinal fasciculus and left optic radiation (**Figure 1, Table 2**).

**Figure 1.**
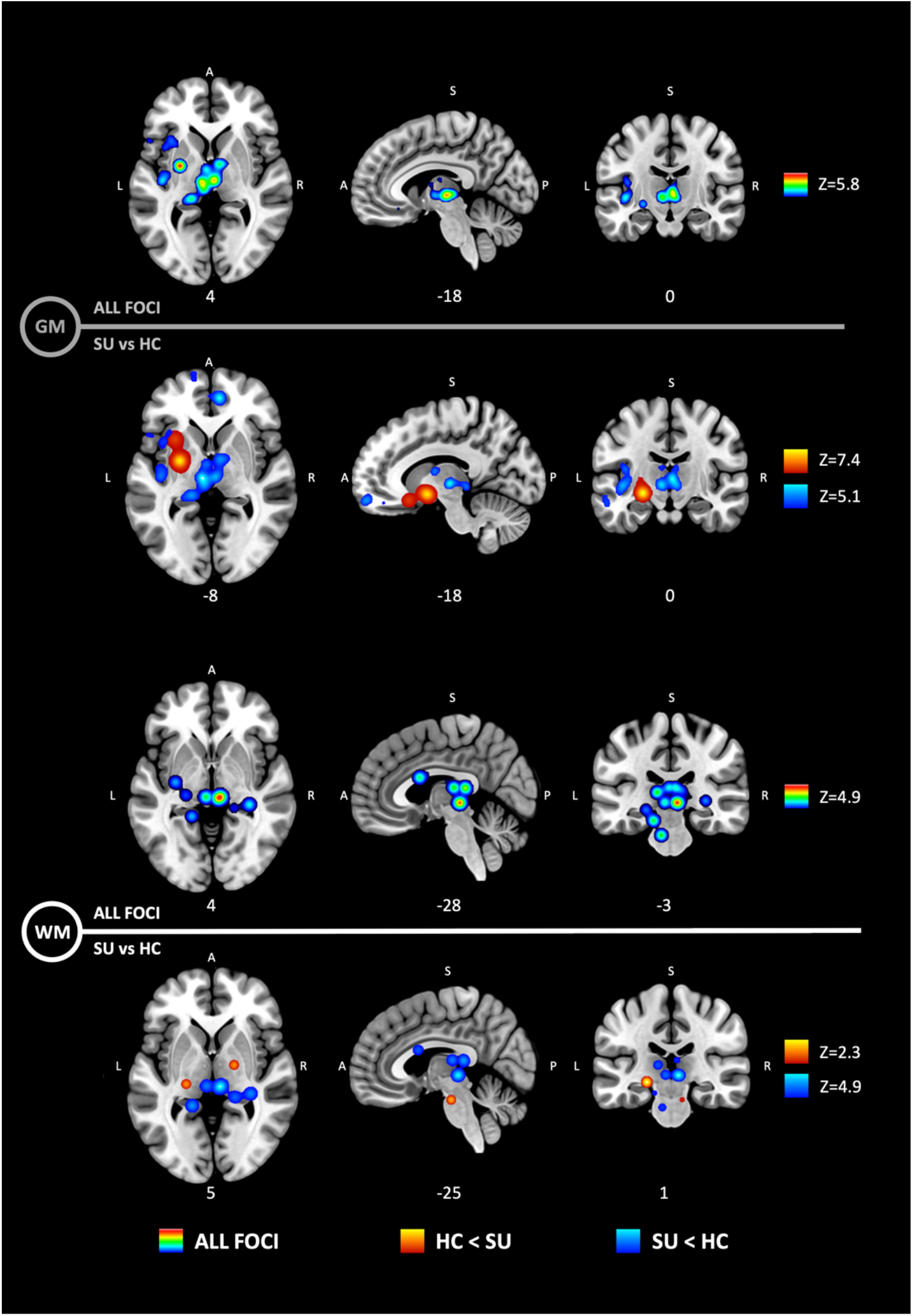
Anatomic likelihood estimation meta-analytic results for studies comparing brain morphological changes between SU and HC, at cluster level inference p < 0.05 (FWE). The primary outcome included GM (top) and WM (bottom) volumetric alterations in SUDs. HC < SU = higher volume with use; SU < HC = lower volume with use. Significant ALE maps show lower volume in thalamus, insula and anterior cingulate cortex in GM, and thalamic radiations in WM; and higher volume in putamen GM, and corticospinal WM tract. Such results support the idea that the entire limbic loop of the basal ganglia shows neuroadaptations produced by SUD. Z, maximum Z-value.

**Table 2.**
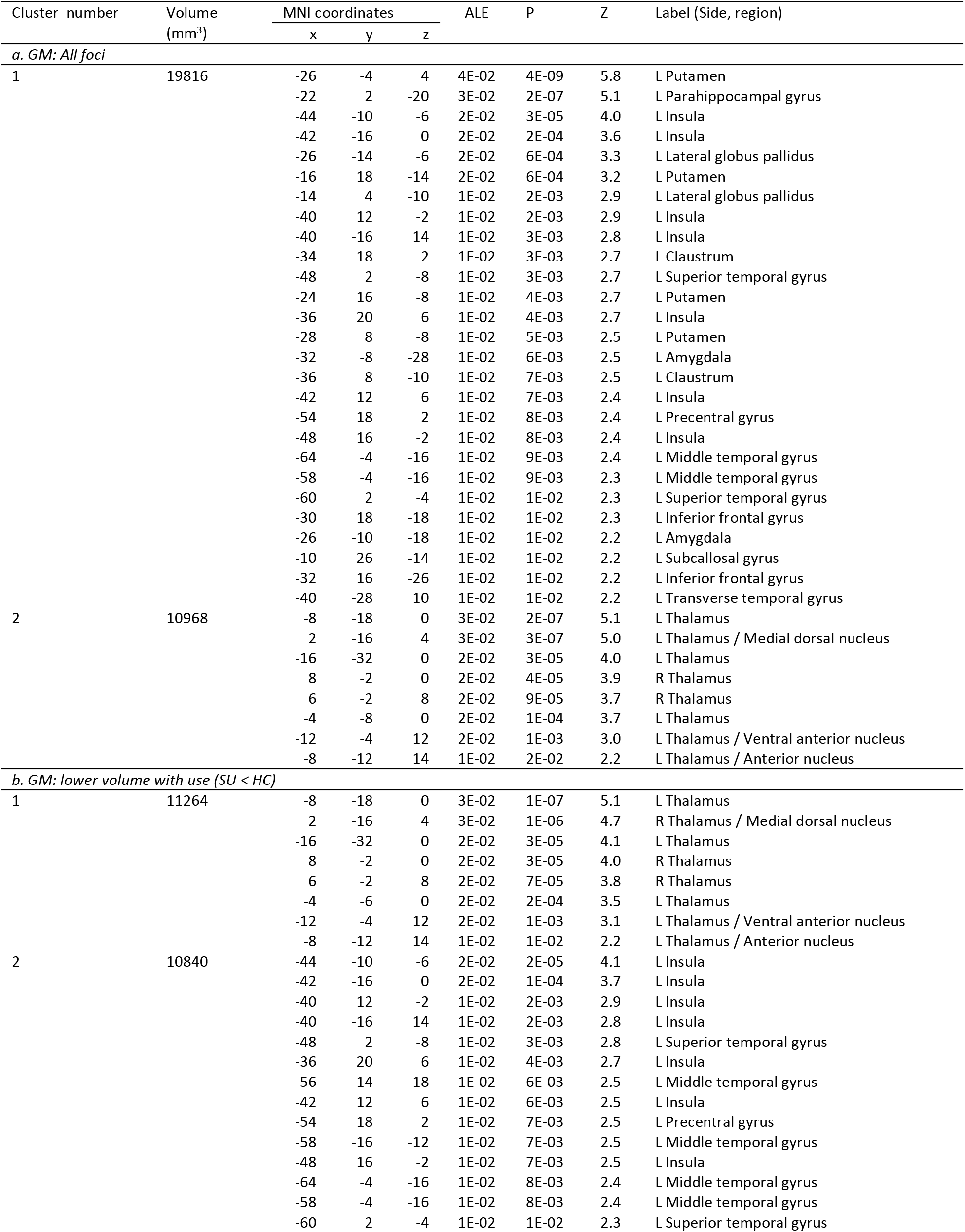
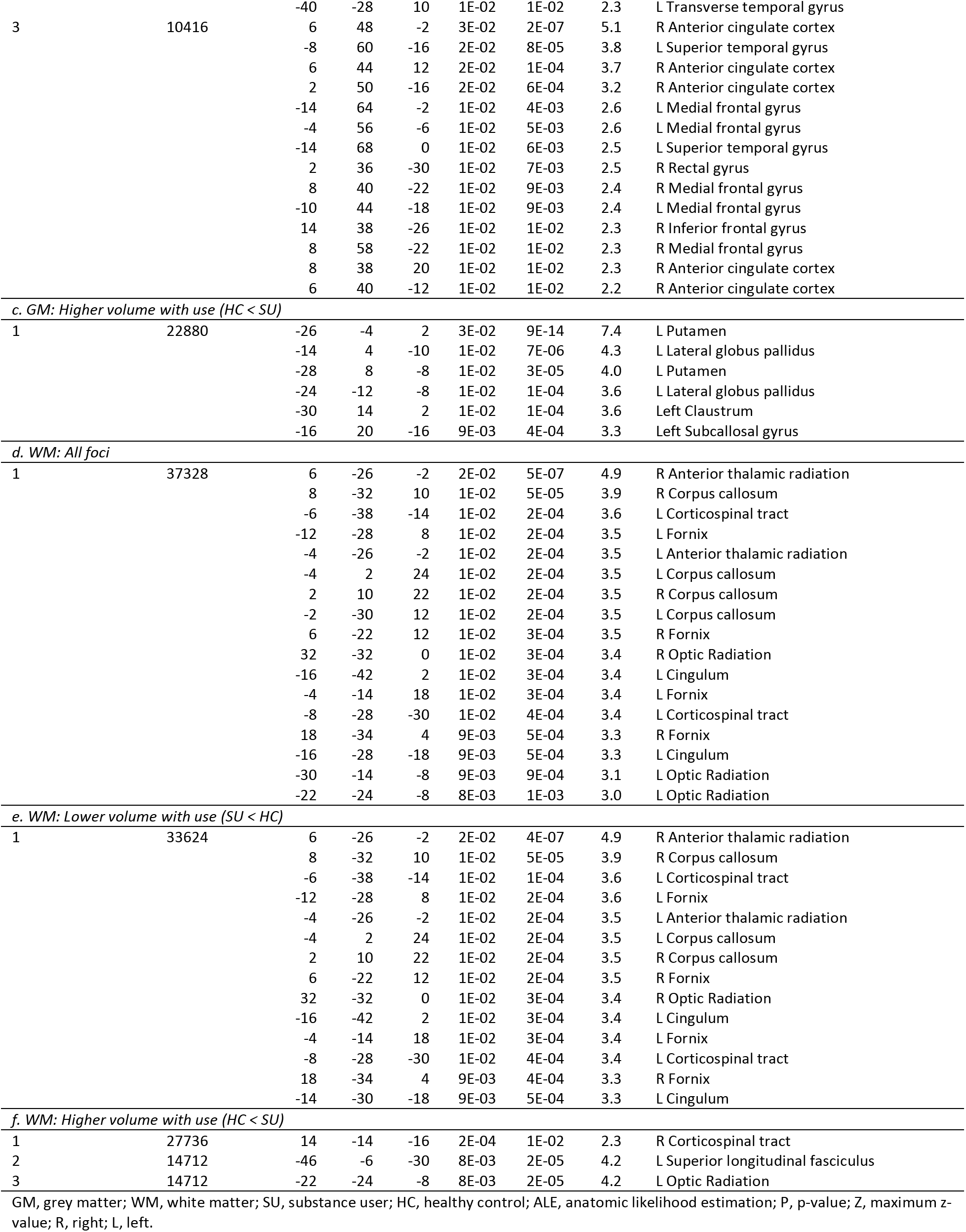
Anatomic likelihood estimation meta-analytic results for studies comparing brain morphological changes between SU and HC, at cluster level inference p < 0.05 (FWE).

#### 3.3.2. Subgroup analyses

*Pre-hoc* subgroup analyses included (1) age of substance user: adolescents vs adults; (2) consumption severity: addiction vs long-term use vs occasional use; (3) type of substance: alcohol vs tobacco vs cannabis vs cocaine vs stimulants vs opioids vs ketamine, and papers that pooled together substances which we termed polysubstance; and (4) associated comorbidities: single vs multiple. Age and comorbidity subgroups resulted in insufficient experiments (foci) to conduct an ALE analysis (<15). However, we found significant ALE maps in the subgroups consumption severity and type of substance (**Figure 2**).

**Figure 2.**
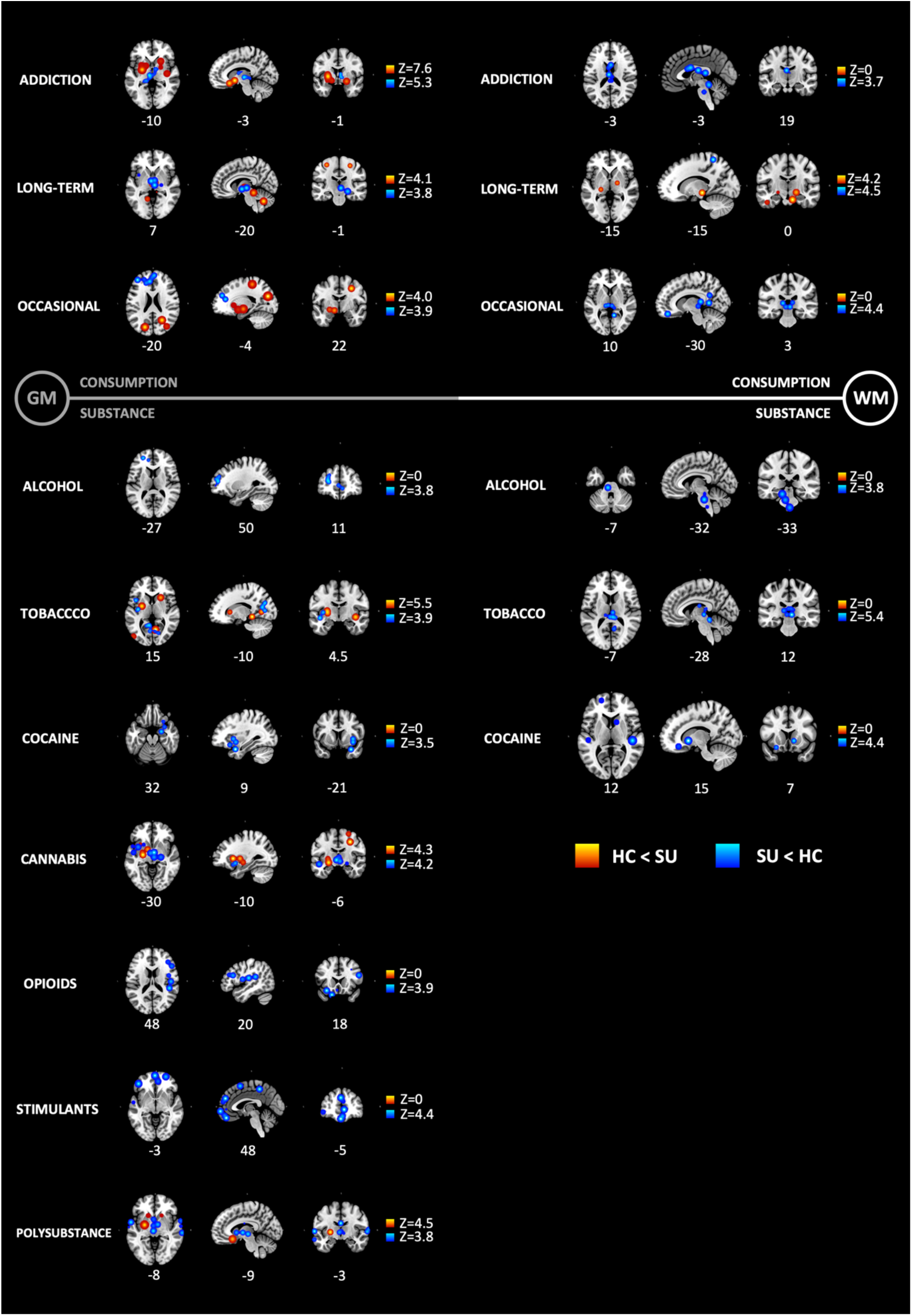
Subgroup anatomic likelihood estimation metaanalytic results for studies comparing brain morphological changes between SU and HC, at cluster level inference p < 0.05 (FWE). Subgroup analyses included consumption severity (top) and type of substance (bottom). HC < SU = higher volume with use; SU < HC = lower volume with use. Significant ALE maps showing lower GM and WM volumes across all types of consumption; higher GM volumes were also shown across all types of consumption; higher WM only in long-term use; lower GM volume in all substances, and higher GM volume only in tobacco, cannabis and polysubstance; lower WM volume in alcohol, tobacco and cocaine, and found no higher WM volume in any substance. Consumption: addiction (k=40) vs long-term use (k=4) vs occasional use (k=6). Substance: alcohol (k=10) vs tobacco (k=11) vs cannabis (k=6) vs cocaine (k=6) vs stimulants (k=3) vs opioids (k=6) vs ketamine(k=1), and papers that pooled together substances which we termed polysubstance (k=7). Z, maximum Z-value.

##### Subgroup analysis by type consumption

The first subgroup meta-analysis reported ALE maps of substance users (SU) against healthy controls (HC), by type of consumption severity (addiction vs long-term use vs occasional use). We found significant ALE maps showing lower GM and WM volumes across all types of consumption. Additionally, higher GM volumes were also shown across all types of consumption, and higher WM only in long-term use (**Figure 2, Supplementary Table 3**).

We conducted contrast analyses between the ALE maps of each subgroup, to determine similarity (conjunction) and/or difference (subtraction) of affected brain regions between the types of consumption (**Figure 3, Supplementary Table 4**). By conducting a contrast analysis of the subgroup ALE maps, we specifically compared and contrasted brain regions across types of consumption severity. Addiction and long-term use were both associated with lower GM volume of the thalamus but differ in terms of lower GM of red nucleus, substantia nigra and putamen. These results support the idea that the thalamus is affected across all levels of SUD severity, and future research should focus on the correlation between SUD progression and the volume/form of the thalamus, as its morphology may predict severity of the disease, and/or monitor the efficacy of treatments and therapies. Addiction and occasional use both show higher volume of the globus pallidus, while differ in lower volume of fronto-temporal areas including the medial frontal gyrus, anterior cingulate cortex and superior temporal gyrus, supporting cortical alterations in occasional use. Finally, long-term use and occasional use share higher volume of somatomotor cortices, due to possible drug intoxication. In terms of WM, addiction and longterm use share lower volume of the anterior thalamic radiations and the corpus callosum, suggesting also a probable correlation between the progression of SUD and the severity in WM structural alteration.

**Figure 3.**
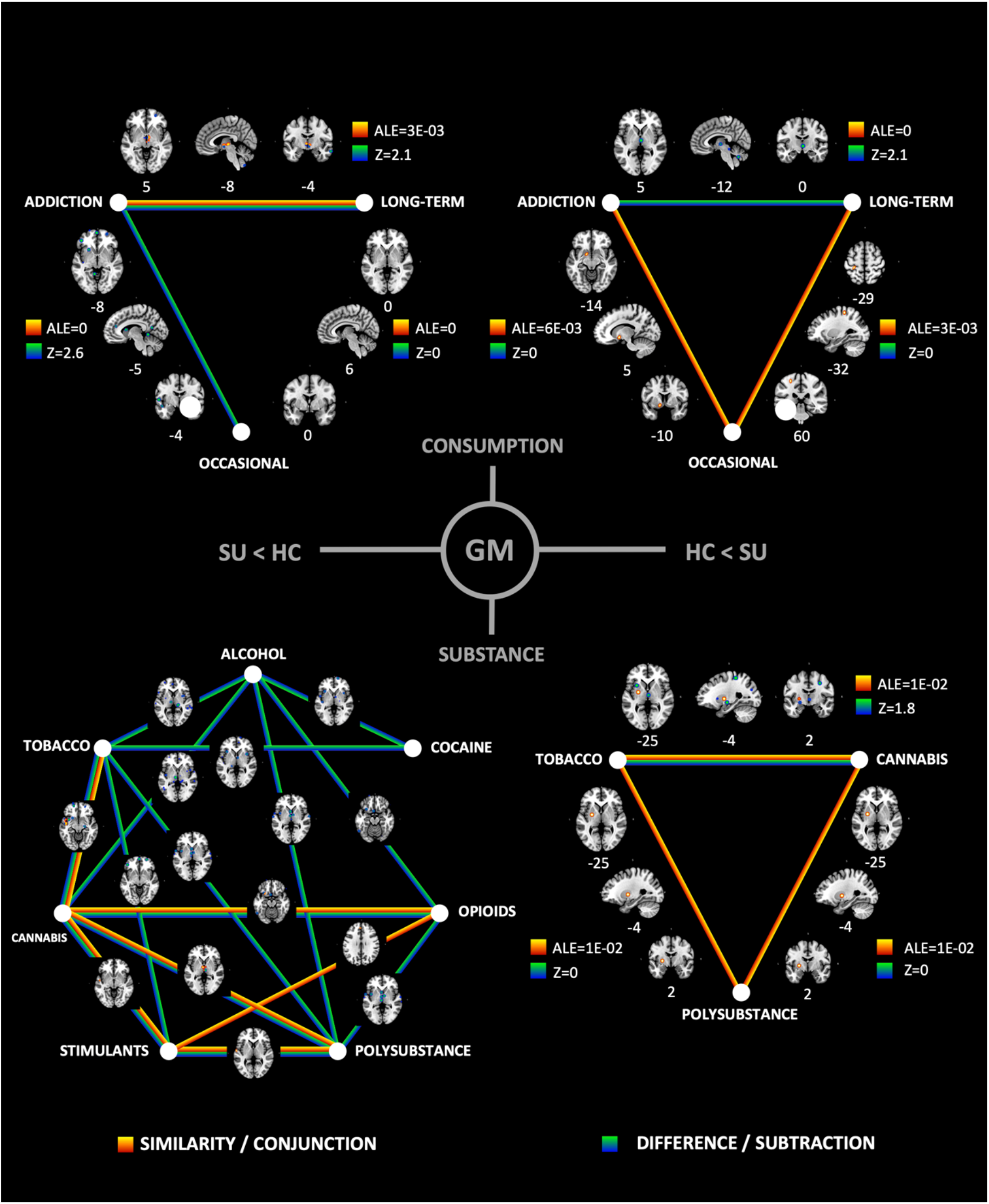
Contrast analyses of subgroup anatomic likelihood estimation meta-analytic results for studies comparing brain morphological changes between SU and HC, at cluster level inference p < 0.05 (FWE). Contrast analyses were performed for consumption severity (top) and type of substance (bottom) subgroups. Subgroups were tested for similarity (conjunction) and difference (subtraction) in a contrast analysis, to illustrate common and/or distinct areas between the elements of each subgroup analysis. ALE, anatomic likelihood estimation value; Z, maximum Z-value.

##### Subgroup analysis by type of substance

In the second subgroup meta-analysis of GM morphometry, we reported ALE maps of substance users (SU) against healthy controls (HC) by type of substance. Given that we only included one publication on ketamine, this substance was not included in the subgroup analysis. We found significant ALE maps showing lower GM volume in all substances, and higher GM volume only in tobacco, cannabis and polysubstance. Also, we found lower WM volume in alcohol, tobacco and cocaine, and found no higher WM volume in any substance (**Figure 2, Supplementary Table 5)**.

We conducted contrast analyses between the ALE maps of each subgroup, to determine similarity (conjunction) and/or difference (subtraction) of affected brain regions between the types of substance (**Figure 3, Supplementary Table 6**). By conducting a contrast analysis of the subgroup ALE maps, we wished to test specifically which brain areas are similar and different between the types of substances. Alcohol, overall, differed with most of the other substances including tobacco, cocaine, cannabis and opioids. Conversely, cannabis shared affected areas with tobacco, opioids, stimulants, and polysubstance. Consistent affected shared areas included thalamus, insula, inferior frontal gyrus and superior temporal gyrus in GM; and anterior thalamic radiation in WM. Although most addictive substances share a common neurobiological process in the reward circuitry, it is evident that neuroadaptations in SUD depend on the type of substance used. Results of this subgroup analysis by substance is valuable for future research into the best approach for therapeutics (pharmacological and behavioral), as treatment effects can be correlated with brain morphometry.

## 4. Discussion

In this systematic review and meta-analysis, we used coordinate-based anatomic likelihood estimation (ALE) to pool the effects of substance use disorders (SUDs) on brain regional volume. We found that the most converging regions with volume pathology in SUDs were putamen, thalamus, insula and anterior cingulate cortex in gray matter (GM), and the thalamic radiation, corticospinal tract and corpus callosum in white matter (WM). We found that consumption severity and type of substance subgroups resulted in significant ALE maps with both shared and distinctive regions involved, supporting converging and divergent effects depending on severity and type of substance use.

### Characteristics of the included studies

Overall, the included publications succeeded in clearly stating their research question, population, inclusion and exclusion criteria, measurements and outcomes. We found that most of the publications failed to report the type of evaluator (e.g. psychiatrist), and some did not mention if the DSM or other tool was used to diagnose SUD. In terms of MRI characteristics and quality of the studies, we found that all included studies used state-of-the-art techniques and statistical tools, and therefore support the standardization of neuroimaging studies as a key element in future research and reproducibility efforts^32–34^. However, a larger effort is needed to provide diagnosis criteria, which would result in improved classifications for future reviews and meta-analyses.

### Primary outcome: altered brain morphometry in SUDs

SUDs seems to disrupt the normal function of the limbic loop of the basal ganglia^3^. Repeated dopaminergic stimulation secondary to substance use, induces persistent neuroplastic adaptations in cortical and subcortical regions that seem to progress with the severity of the SUD^42^. Indeed, we found volumetric alterations in putamen, thalamus, insula and anterior cingulate cortex in GM, and internal capsule and thalamic radiations in WM, supporting the idea that the entire limbic loop of the basal ganglia shows neuroadaptations produced by SUDs.

Higher volume of the putamen may be explained by the repeated glutamatergic spikes onto dopamine neurons (VTA/SNc) and into MSN in dorsal and ventral striatum, due to repeated substance use, and supported by behavioral changes in reward responsivity and habituation that characterize SUDs. Noticeably, almost all regions of the neocortex project direct input to the striatum. Most of these projections come from association areas in frontal and parietal lobes, with contributions from temporal, insular, and cingulate cortices. These projections (corticostriatal pathway) travel via the internal capsule to reach the caudate and putamen^43^. We also found higher WM volume of the internal capsule in SUDs, suggesting neuroadaptive processes in this pathway as well. It has been suggested that SUDs or addiction are a disease of self-control^44^. Although the study of SUDs has been focused mainly on the role of dopamine and the reward system, new findings of clinical studies have revealed neuroplastic mechanisms in frontocortical regions that may underlie reward-seeking behavior^14^. In susceptible individuals, certain stimuli may activate strong urges that are not congruent with a given context. The lack of a proper inhibitory control may keep these urges in control up to a point, when stronger impulses and deficient inhibition result in impulsive or compulsive behavior^45^. Current models of SUDs suggest that impulsivity and compulsivity characterize the pathological behavior and help explain our structural results^3^.

It has been proposed that the insula and the anterior cingulate cortex form the salience network (SN), that coordinates between the default mode network (DMN) and the central executive network (CEN)^46^. In our study we found lower volume of the insula, a region whose morphology has been associated with substance use compulsion and severity^47^. The insula plays a major role in interoception by integrating information from the internal physiological state, and projecting information to the ACC, ventral striatum and prefrontal cortex to initiate adaptive responses^48^. In SUDs, the insula’s ability to switch between networks seems to be affected, as well as its functional connectivity with the ACC, amygdala and putamen^49–51^. Similarly, SUD neuroimaging studies have shown disrupted activity of the ACC^3^, involved in inhibitory control^52^, and altered connectivity with the insula^99^. The rostral part of the ACC is implicated in error-related responses, including affective processing, and the caudal part of the ACC is associated with detection of conflict to recruit cognitive control^53^. Thus, reduction in inputs from prefrontal and cingulate cortices into striatum may disrupt the control over action selection^54^.

Finally, we found that SUDs were associated with lower thalamic GM/WM across several substances including alcohol, cocaine, nicotine, methamphetamine, opioids and cannabis^55,56^. Reduced structural and functional integrity of the thalamus and its connectivity appear to be associated with the severity of SUD^57^. Overall, there are brain regions consistently affected in all SUDs, with diverging MRI manifestations (higher vs lower volume) suggesting different underlying structural pathology between brain regions.

### Common and distinct patterns of brain volume alterations across consumption severity

We found that the effect of substance use in the brain seems to vary across the severity of consumption. Cortical structures seem affected in occasional use, while established addictive consumption (addiction) seems to also affect subcortical regions of the brain such as thalamus and basal ganglia. In addiction, such disrupted GM areas may presumably be co-affected with its respective WM thalamic radiation and corpus callosum connection, as seen in our results. We also found that occasional use affects WM tracts of the cingulum, connecting the limbic system with areas such as the cingulate gyrus, entorhinal cortex and temporal lobe. Neuroimaging studies have found that disruption of the posterior cingulum is associated to cognitive impairment^58^. The forceps minor connects the lateral and medial surfaces of the frontal lobes and crosses the midline via the genu of the corpus callosum^43^, and also showed structural alterations in occasional use. Along with the anterior thalamic radiation, the forceps minor connects ACC and striatum to the anterior frontal regions, modulating executive functions^59^.

Various physiological mechanisms such as oxidative stress, mitochondrial dysfunction or neurotrophic factor dysfunction might account for the observed cortical GM volume reductions in occasional use^60^. However, repeated dopaminergic stimulation from substance abuse produce neuroadaptations (e.g., dendritic morphology and ionotropic glutamate receptors), that result in long-term potentiation (LTP) and long-term depression (LTD)^12^ of the basal ganglia neurocircuitry^3^. These results suggest that cortical morphological pathology in SUDs appears before subcortical pathology or that subcortical pathology is only seen when addiction is established. This needs to be explored further with longitudinal studies.

### Common and distinct patterns of brain volume alterations across types of substances

Reward processes are shared between substances, namely repeated stimulation into the VTA which releases dopamine into the ventral striatum^3^. However, the stimulation of the mesolimbic system depends on the different molecular targets for each kind of substance. For example, alcohol, unlike most other drugs, affects a wide range of targets and indirectly increases dopamine in the NAc^61^. Stimulants like amphetamine and cocaine block dopamine transporters, thus increasing dopamine in NAc^62^. Cannabis activates receptors that release neurotransmitters (GABA/Glutamate), modulating the activity of the mesolimbic system. Opioids, agonists of mu opioid receptors (MOR) in VTA, increase striatal dopamine release^63^. Nicotine and its interactions with nicotinic acetylcholine receptors, increases neuronal activity in VTA^64^. In our results, most of the substances show a convergent effect and region, namely lower volume of the thalamus.

Divergently, alcohol seems to affect frontal areas including superior and medial frontal gyrus, as well as ACC. Tobacco use shows a myriad of alterations including lower volume in insula and posterior areas of the DMN, such as PCC and precuneus. Cocaine users show lower volume of the claustrum, a structure that connects prefrontal areas with the thalamus, and has close proximity to the insula and putamen^65^. Cannabis use reduces the volume of temporal areas and thalamus, and increases the volume of putamen, while opioid use affects cortical fronto-temporal areas. Stimulant use mainly reduces GM volume of the frontal lobe. Polysubstance studies, as expected, show a wide variety of affected areas including lower volume of the anterior cingulate gyrus, thalamus, and superior temporal gyrus, and show higher volume of the subcallosal gyrus. In terms of WM, the affected convergent regions were the corticospinal tract, anterior thalamic radiation, the corpus callosum, and the cingulum. Overall, different substances show convergent and divergent morphological pathology, suggesting different physiopathology and possibly therapeutic approaches in SUDs that need to be considered.

### Limitations

To conduct the anatomic likelihood estimation meta-analysis, we pooled peak coordinates derived from the included studies, rather the original raw structural MRI images. The accuracy of our findings relies on the result of a statistical estimation of coordinate-based anatomic foci (input), treated as spatial probability distributions centered at the given coordinates. The heterogeneity among the methods used in the included studies, such as preprocessing software, smoothing, statistical thresholds, characteristics of the participants, medication history and comorbidity, represent potential confounders. Meta-regression analysis is not compatible with GingerALE, which would have shown important insights when testing for heterogeneity (e.g. age of participants, age of first use, total years of SUD).

As traditional meta-analyses, coordinate-based meta-analyses such as ALE can be subject to different forms of publication bias which may impact results and invalidate findings (e.g., the file drawer problem). We performed the Fail-Safe N analysis (FSN)^39^ as a measure of robustness against potential publication bias. It is estimated for normal human brain mapping that a 95% confidence interval for the number of studies that report no local maxima varies from 5 to 30 per 100 published studies. Using the upper bound and the fact that our meta-analysis consists of 50 experiments, a possible estimate for the number of experiments that remain in the file drawer is 15. Therefore, the minimum FSN was defined as 15 (**Supplementary Table 7**). In our study, we tested 11 clusters resulting from our primary outcomes. We found that all clusters showed an FNR greater or equal than the minimum imposed of 15. FNR was >350 for clusters resulting from all GM foci analysis; and >300 for clusters resulting from all WM foci analysis. Thus, indicating a robust convergence of foci in these regions but also indicating that proportionally fewer studies are needed to obtain this effect. Two clusters from the comparison GM SU<HC showed FSN than laid between the lower and upper boundary. One cluster showed exactly the lower boundary.

In our review, studies are not really investigating long-term measurements necessary to show that SUD is causing decrease or increase of brain tissue, as a longitudinal design might provide; but they rather examine brain morphometry in established substance user compared to nonusers. Socio-economic and educational background data on participants are lacking in most of the studies, limiting the potential for statistical correction using naturalistic environmental confounders. In the consumption and substance subgroup analyses, the number of experiments for each category of the subgroup analyses was unmatched (e.g., addiction 59%, occasional use 33% and long-term use 8%). Although the ALE method weights the result on the number of participants per experiment, the resulting ALE maps of subgroup and contrast analyses should be interpreted with caution.

The progression from initial drug use to established SUD may depend on age and developmental stage^66^. Critical periods of development are characterized by functional neuroplastic mechanisms that may be easily altered by pathological neuroadaptations due to SUD^5^. For example, delays in maturation associated with drug exposure, genetics, or social environment, may increase risky behaviors in adolescents^67^. Brain imaging studies have found altered structure of prefrontal cortices associated with higher risk for SUD in adolescents^68^, suggesting that control executive functions such as decision making and impulse control (inhibition) are immature^69^. Unfortunately, the neurobiological underpinnings of neuroadaptations for both functional development and SUD, are not fully understood, in part, by a high variability in VBM results^70^. In this review, the included studies failed to report enough experiments (foci<15), to conduct an age subgroup analysis (e.g., adolescents vs adults).

SUDs are frequently co-diagnosed with psychiatric and neurological disorders^71^. For example, research suggests that adolescents with SUD have high rates of co-occurring mental illness, up to 60%^72^. The most common psychiatric comorbidities with SUD include anxiety disorders, post-traumatic stress disorder, depression, bipolar disorder, attention-deficit hyperactivity disorder, psychosis, borderline personality disorder, and schizophrenia. Noticeably, establishing causality or directionality between mental illness and SUD is difficult, however, common risk factors are shared^73^. Additionally, recent research has focused on the neurological effects of SUD, rather than as comorbid, co-occurring alterations^74^ (e.g., SUD and Parkinson’s disease). In this review, the included studies failed to report enough experiments (foci<15), to conduct a comorbidity subgroup analysis (e.g., pure addiction vs comorbid addiction). Nevertheless, it is important to recognize that mental illness and SUD share alterations in the same neurotransmitter systems (e.g., dopaminergic^4^) and in brain areas involved in reward, decision making, impulse control and emotion^75^.

## 5. Conclusions

In conclusion, the present systematic review and meta-analysis of voxel-based morphometry neuroimaging studies provides evidence of common and distinct morphological gray matter and white matter pathology in substance use disorders. We found consistent morphometric alterations in regions of the insula, anterior cingulate cortex, basal ganglia (putamen), and thalamus, with their respective white matter thalamic radiation and internal capsule bundle. Our subgroup analysis showed distinct volume alterations depending on the type of consumption (occasional vs long-term vs addiction) and type of substance. This evidence may help future studies to better understand substance use disorders and possible new therapeutic approaches.

## Acknowledgments

Scholarship for Sofia Fernandez-Lozano funded by CONACYT No. 476284. This study was supported by public fund CONACYT FOSISS No. 0260971

## Conflict of interest

There is no actual or potential financial and other conflict of interest related to this manuscript.

Supplementary information (SI) is available at MP’s website.

## SUPPLEMENTARY INFORMATION

### METHODS

#### Primary literature search and selection

This systematic review and meta-analysis followed procedures from the Cochrane Handbook for Systematic Reviews^35^, and from the Center for Reviews and Dissemination (https://www.york.ac.uk/crd/). The review protocol was pre-registered in PROSPERO (<u>CRD42017071222</u>). The PRISMA statement was used to address relevant items in this systematic review and meta-analysis^36^.

#### Search strategy

We conducted a systematic literature search in PubMed, Scopus and PsycInfo, using both keywords and MeSH terms for articles published to the end of July 2018. Keyword terms and MeSH terms included: substance-related disorders, alcohol-related disorders, amphetamine-related disorders, cocaine-related disorders, inhalant abuse, marijuana abuse, substance abuse (intravenous), tobacco use disorder, drug utilization, cannabis, cocaine, crack cocaine, alcoholics, amphetamine, methamphetamine, N-Methyl-3,4-methylenedioxyamphetamine. No restrictions were placed on study design, but in order to be eligible for inclusion, the studies must have used VBM analyses (search terms in Appendix).

#### Study eligibility

Studies were included if they met the following criteria: (1) an original report of whole-brain VBM analyses, (2) the study population included substance users above 18 years of age, with a continuous level of usage (at least once a month in the last 6 months), (3) studies included a substance user group and a non-exposed group (healthy control group), and (4) results were reported in stereotactic coordinates either Talairach or Montreal Neurological Institute (MNI) three-dimensional-coordinate system. If any of these data points were not reported in the paper, we contacted authors to retrieve this information. We contacted a total of 4 authors, with no response.

Studies were excluded using the following criteria: (1) review articles with no original experimental data, (2) neuroimaging data from non-MRI studies (e.g. PET), (3) the type of addiction was gambling, internet, etc. (non-substance addiction), and (4) the VBM analyses were region-of-interest (ROI)-based and not whole-brain-based.

Two reviewers (VP and ST) independently screened by title and abstract and selected articles for full-text review, and also performed full-text reviews. Screening and data extraction were performed using the Covidence tool^37^. Any disagreements that arose between the reviewers were resolved through discussion or by a third and/or fourth reviewer (SA / EGV). A total of 23 disagreements were resolved in the title and abstract screening, and 14 disagreements were resolved during full-text screening. A total of 50 studies fulfilled criteria and were therefore included for data extraction.

#### Data extraction

From each study, the following variables were extracted: first author, year of publication, population of interest, use of the Diagnostic and Statistical Manual of Mental Disorders (DSM), type of addiction (pure or dual = comorbidity), number of participants, sex, age, type of substance, type of consumption, years of consumption, age of onset of consumption, MRI-system, MRI-model, Head coil, image acquisition, T1w sequence, voxel size and analysis software. The main outcome to extract was any change in gray and/or white matter analyzed using VBM, in stereotactic coordinates, comparing a substance user group and a healthy control group.

#### Search strings

Last search date: 18.07.2018

#### PsycInfo

(voxel-based morphometry OR voxel based morphometry OR vbm) AND (addiction OR substance-related disorders OR alcohol-related disorders OR amphetamine-related disorders OR cocaine-related disorders OR inhalant abuse OR marijuana abuse OR substance abuse intravenous OR tobacco use disorder OR drug utilization OR drug abuse OR drug dependency OR substance utilization OR substance abuse OR substance dependency OR cannabis OR marihuana OR marijuana OR tetrahydrocannabinol OR thc OR cocaine OR crack cocaine OR alcohol OR alcoholics OR amphetamine OR methamphetamine OR n-methyl-3,4-methylenedioxyamphetamine OR mdma OR ecstasy OR heroin OR nicotine OR tobacco) 249 results

#### Scopus

<u>((TITLE-ABS-KEY (**voxel-based morphometry**) OR TITLE-ABS-KEY (**voxel based morphometry**) OR TITLE-ABS-KEY (**vbm**)) AND (TITLE-ABS-KEY (**addiction**) OR TITLE-ABS-KEY (**substance-related disorders**) OR TITLE-ABS-KEY (**alcohol-related disorders**) OR TITLE-ABS-KEY (**amphetamine-related disorders**) OR TITLE-ABS-KEY (**cocaine-related disorders**) OR TITLE-ABS-KEY (**inhalant abuse**) OR TITLE-ABS-KEY (**marijuana abuse**) OR TITLE-ABS-KEY (**substance abuse intravenous**) OR TITLE-ABS-KEY (**tobacco use disorder**) OR TITLE-ABS-KEY (**drug utilization**) OR TITLE-ABS-KEY (**drug abuse**) OR TITLE-ABS-KEY (**drug dependency**) OR TITLE-ABS-KEY (**substance utilization**) OR TITLE-ABS-KEY (**substance abuse**) OR TITLE-ABS-KEY (**substance dependency**) OR TITLE-ABS-KEY (**cannabis**) OR TITLE-ABS-KEY (**marihuana**) OR TITLE-ABS-KEY (**marijuana**) OR TITLE-ABS-KEY (**tetrahydrocannabinol**) OR TITLE-ABS-KEY (**thc**) OR TITLE-ABS-KEY (**cocaine**) OR TITLE-ABS-KEY (**crack cocaine**) OR TITLE-ABS-KEY (**alcohol**) OR TITLE-ABS-KEY (**alcoholics**) OR TITLE-ABS-KEY (**amphetamine**) OR TITLE-ABS-KEY (**methamphetamine**) OR TITLE-ABS-KEY (**n-methyl-3,4-methylenedioxyamphetamine**) OR TITLE-ABS-KEY (**MDMA**) OR TITLE-ABS-KEY (**ecstasy**) OR TITLE-ABS-KEY (**heroin**) OR TITLE-ABS-KEY (**nicotine**) OR TITLE-ABS-KEY (**tobacco**)))</u> 327 results

#### PubMed

(((((((“voxel-based morphometry”[All Fields]) ((“voxel-based”[All Fields]) AND (“morphometry”[All Fields])) OR (“voxel based morphometry”[All Fields]) OR ((“voxel”[All Fields]) AND (“based”[All Fields]) AND (“morphometry”[All Fields])) OR (“vbm”[All Fields])))) AND ((((“addiction”[All Fields]) OR (“substance-related disorders”[MeSH Terms]) OR ((“substance-related”[All Fields]) AND (“disorders”[All Fields])) OR (“substance-related disorders”[All Fields]) OR (“substance related disorders”[All Fields]) OR ((“substance”[All Fields]) AND (“related”[All Fields]) AND (“disorders”[All Fields])) OR (“alcohol-related disorders”[MeSH Terms]) OR (“alcohol-related disorders”[All Fields]) OR (“alcohol related disorders”[All Fields]) OR ((“alcohol-related”[All Fields]) AND (“disorders”[All Fields])) OR ((“alcohol”[All Fields]) AND (“related” [All Fields]) AND (“disorders” [All Fields])) OR (“amphetamine-related disorders”[MeSH Terms]) OR (“amphetamine-related disorders”[All Fields]) OR ((“amphetamine-related” [All Fields]) AND (“disorders” [All Fields])) OR (“amphetamine related disorders” [All Fields]) OR ((“amphetamine” [All Fields]) AND (“related” [All Fields]) AND (“disorders” [All Fields])) OR (“cocaine-related disorders”[MeSH Terms]) OR (“cocaine-related disorders”[All Fields]) OR ((“cocaine-related” [All Fields]) AND (“disorders” [All Fields])) OR (“cocaine related disorders” [All Fields]) OR ((“cocaine” [All Fields]) AND (“related” [All Fields]) AND (“disorders” [All Fields])) OR (“inhalant abuse”[MeSH Terms]) OR (“inhalant abuse”[All Fields]) OR ((“inhalant” [All Fields]) AND (“abuse” [All Fields])) OR (“marijuana abuse”[MeSH Terms]) OR (“marijuana abuse”[All Fields]) OR ((“marijuana” [All Fields]) AND (“abuse” [All Fields])) OR (“substance abuse intravenous”[MeSH Terms]) OR (“substance abuse intravenous”[All Fields]) OR ((“substance” [All Fields]) AND (“abuse” [All Fields]) AND (“intravenous” [All Fields])) OR (“tobacco use disorder”[MeSH Terms]) OR (“tobacco use disorder”[All Fields]) OR ((“tobacco” [All Fields]) AND (“use” [All Fields]) AND (“disorder” [All Fields])) OR (“drug utilization”[MeSH Terms]) OR (“drug utilization”[All Fields]) OR ((“drug” [All Fields]) AND (“utilization” [All Fields])) OR (“drug abuse”[All Fields]) OR ((“drug” [All Fields]) AND (“abuse” [All Fields])) OR (“drug dependency”[All Fields]) OR ((“drug” [All Fields]) AND (“dependency” [All Fields])) OR (“substance utilization”[All Fields]) OR ((“substance” [All Fields]) AND (“utilization” [All Fields])) OR (“substance abuse”[All Fields]) OR ((“substance” [All Fields]) AND (“abuse” [All Fields])) OR (“substance dependency”[All Fields]) OR ((“substance” [All Fields]) AND (“dependency” [All Fields])) OR (“cannabis”[MeSH Terms]) OR (“cannabis”[All Fields]) OR (“marihuana”[All Fields]) OR (“marijuana”[All Fields]) OR (“tetrahydrocannabinol”[All Fields]) OR (“thc”[All Fields]) OR (“cocaine”[MeSH Terms]) OR (“cocaine”[All Fields]) OR (“crack cocaine”[MeSH Terms]) OR (“crack cocaine”[All Fields]) OR ((“crack” [All Fields]) AND (“cocaine” [All Fields])) OR (“alcohol”[All Fields]) OR (“alcoholics”[MeSH Terms]) OR (“alcoholics”[All Fields]) OR (“amphetamine”[MeSH Terms]) OR (“amphetamine”[All Fields]) OR (“methamphetamine”[MeSH Terms]) OR (“methamphetamine”[All Fields]) OR (“n-methyl-3,4-methylenedioxyamphetamine”[MeSH Terms]) OR (“n-methyl-3,4-methylenedioxyamphetamine”[All Fields]) OR (“mdma”[All Fields]) OR (“ecstasy”[All Fields]) OR (“heroin”[MeSH Terms]) OR (“heroin”[All Fields]) OR (“nicotine”[MeSH Terms]) OR (“nicotine”[All Fields]) OR (“tobacco”[MeSH Terms]) OR (“tobacco”[All Fields]))))

243 results

#### Fail-Safe N analysis (FSN)

Coordinate-based meta-analyses such as ALE can be subject to different forms of publication bias which may impact results and invalidate findings (e.g., the file drawer problem). We performed the Fail-Safe N analysis (FSN) as a measure of robustness against potential publication bias. It refers to the amount of contra-evidence that can be added to a meta-analysis before the results change, and can be obtained for each cluster that survives thresholding in an ALE meta-analysis. A higher FSN indicates more stable results and hence a higher robustness. It is estimated for normal mapping that a 95% confidence interval for the number of studies that report no local maxima varies from 5 to 30 per 100 published studies. Using the upper bound and the fact that our meta-analysis consists of 50 experiments, a possible estimate for the number of experiments that remain in the file drawer is 15. Therefore, the minimum FSN was defined as 15 (Results in **Supplementary Table 7**).

### RESULTS

**Supplementary Figure 1.**
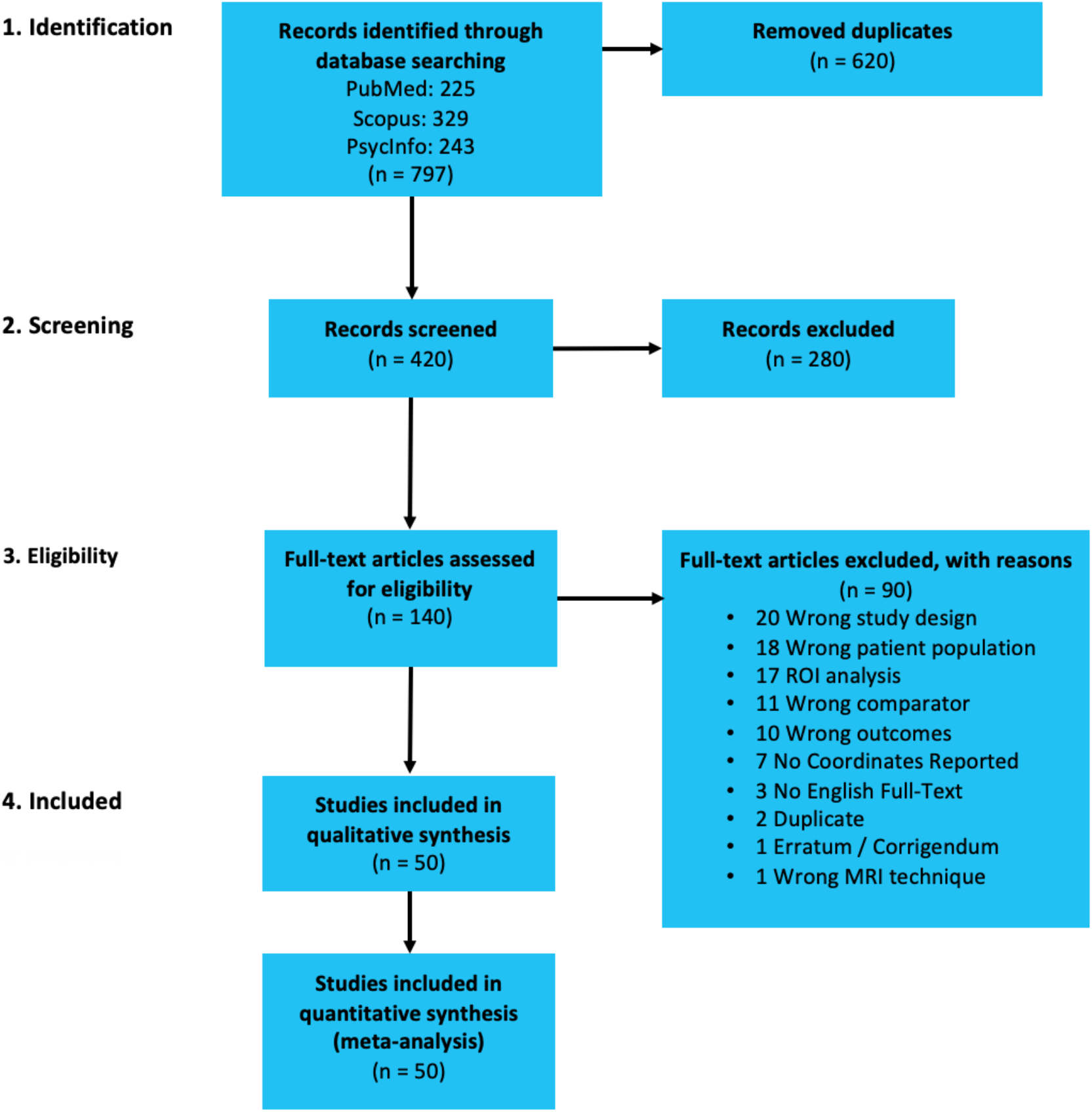
PRISMA flowchart for literature search process.

### PRISMA CHECKLIST

#### SI = Supplementary Information

| Section/topic | # | Checklist item | Reported on page # |
| --- | --- | --- | --- |
| **TITLE** |  |  |  |
| Title | 1 | Identify the report as a systematic review, meta-analysis, or both. | 1 |
| **ABSTRACT** |  |  |  |
| Structured summary | 2 | Provide a structured summary including, as applicable: background; objectives; data sources; study eligibility criteria, participants, and interventions; study appraisal and synthesis methods; results; limitations; conclusions and implications of key findings; systematic review registration number. | 2 |
| **INTRODUCTION** |  |  |  |
| Rationale | 3 | Describe the rationale for the review in the context of what is already known. | 3 |
| Objectives | 4 | Provide an explicit statement of questions being addressed with reference to participants, interventions, comparisons, outcomes, and study design (PICOS). | 4 |
| **METHODS** |  |  |  |
| Protocol and registration | 5 | Indicate if a review protocol exists, if and where it can be accessed (e.g., Web address), and, if available, provide registration information including registration number. | 5 |
| Eligibility criteria | 6 | Specify study characteristics (e.g., PICOS, length of follow-up) and report characteristics (e.g., years considered, language, publication status) used as criteria for eligibility, giving rationale. | 5 & SI |
| Information sources | 7 | Describe all information sources (e.g., databases with dates of coverage, contact with study authors to identify additional studies) in the search and date last searched. | 5 & SI |
| Search | 8 | Present full electronic search strategy for at least one database, including any limits used, such that it could be repeated. | SI |
| Study selection | 9 | State the process for selecting studies (i.e., screening, eligibility, included in systematic review, and, if applicable, included in the meta-analysis). | 5 & SI |
| Data collection process | 10 | Describe method of data extraction from reports (e.g., piloted forms, independently, in duplicate) and any processes for obtaining and confirming data from investigators. | 5 & SI |
| Data items | 11 | List and define all variables for which data were sought (e.g., PICOS, funding sources) and any assumptions and simplifications made. | 5 & SI |
| Risk of bias in individual studies | 12 | Describe methods used for assessing risk of bias of individual studies (including specification of whether this was done at the study or outcome level), and how this information is to be used in any data synthesis. | 5 & SI |
| Summary measures | 13 | State the principal summary measures (e.g., risk ratio, difference in means). | 5 |
| Synthesis of results | 14 | Describe the methods of handling data and combining results of studies, if done, including measures of consistency (e.g., $I^2$) for each meta-analysis. | 5 & 6 |
| Risk of bias across studies | 15 | Specify any assessment of risk of bias that may affect the cumulative evidence (e.g., publication bias, selective reporting within studies). | 6 |
| Additional analyses | 16 | Describe methods of additional analyses (e.g., sensitivity or subgroup analyses, meta-regression), if done, indicating which were pre-specified. | 6 |
| **RESULTS** |  |  |  |
| Study selection | 17 | Give numbers of studies screened, assessed for eligibility, and included in the review, with reasons for exclusions at each stage, ideally with a flow diagram. | 6 & SI |
| Study characteristics | 18 | For each study, present characteristics for which data were extracted (e.g., study size, PICOS, follow-up period) and provide the citations. | 6 |
| Risk of bias within studies | 19 | Present data on risk of bias of each study and, if available, any outcome level assessment (see item 12). | 5, 7 & SI |
| Results of individual studies | 20 | For all outcomes considered (benefits or harms), present, for each study: (a) simple summary data for each intervention group (b) effect estimates and confidence intervals, ideally with a forest plot. | 7 |
| Synthesis of results | 21 | Present results of each meta-analysis done, including confidence intervals and measures of consistency. | 7 |
| Risk of bias across studies | 22 | Present results of any assessment of risk of bias across studies (see Item 15). | SI |
| Additional analysis | 23 | Give results of additional analyses, if done (e.g., sensitivity or subgroup analyses, meta-regression [see Item 16]). | 8 & SI |
| **DISCUSSION** |  |  |  |
| Summary of evidence | 24 | Summarize the main findings including the strength of evidence for each main outcome; consider their relevance to key groups (e.g., healthcare providers, users, and policy makers). | 15 |
| Limitations | 25 | Discuss limitations at study and outcome level (e.g., risk of bias), and at review-level (e.g., incomplete retrieval of identified research, reporting bias). | 18 |
| Conclusions | 26 | Provide a general interpretation of the results in the context of other evidence, and implications for future research. | 20 |
| **FUNDING** |  |  |  |
| Funding | 27 | Describe sources of funding for the systematic review and other support (e.g., supply of data); role of funders for the systematic review. | 20 |

**Supplementary Table 1.**
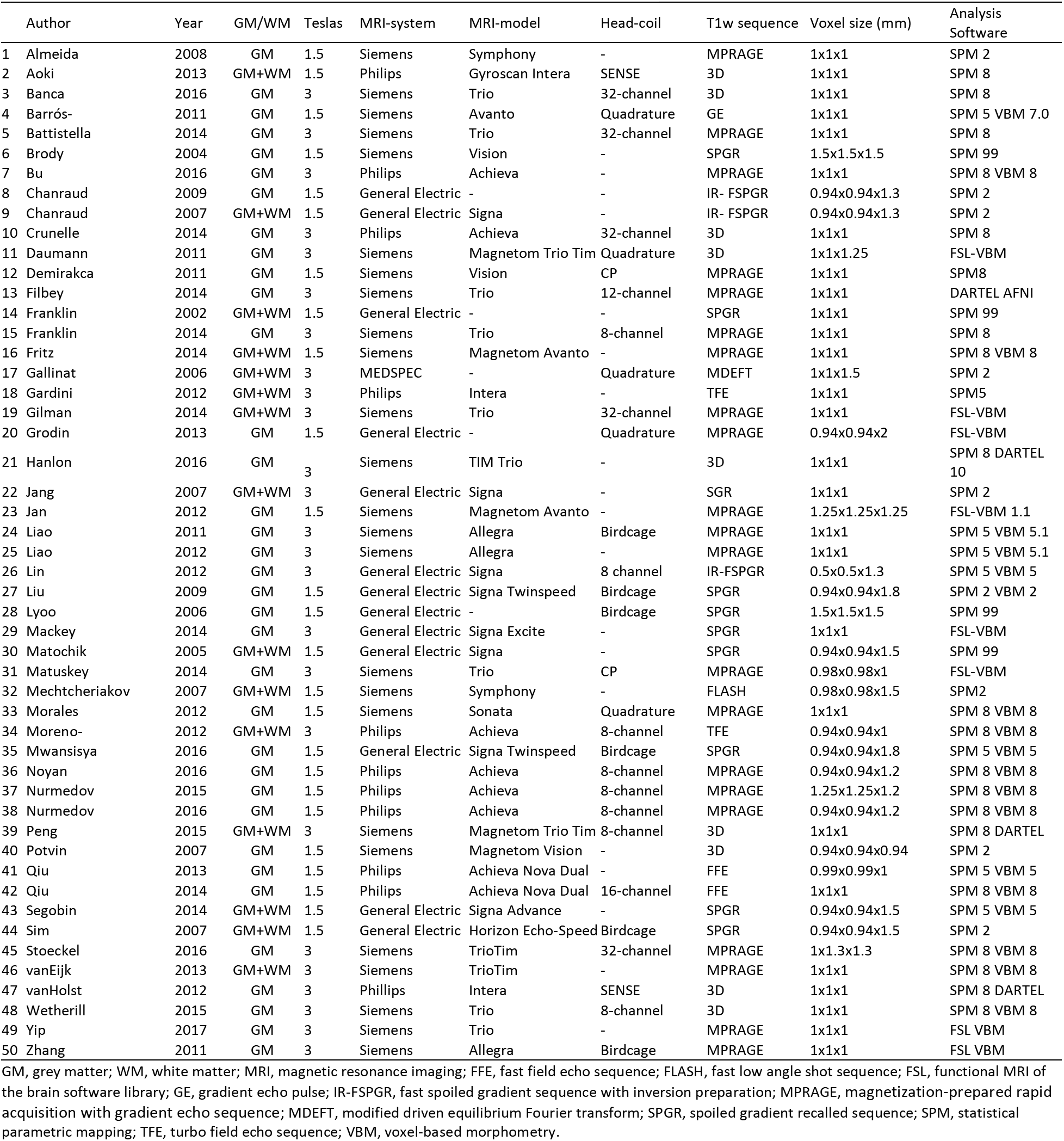
Characteristics of MRI acquisition and analysis.

**Supplementary Table 2.**
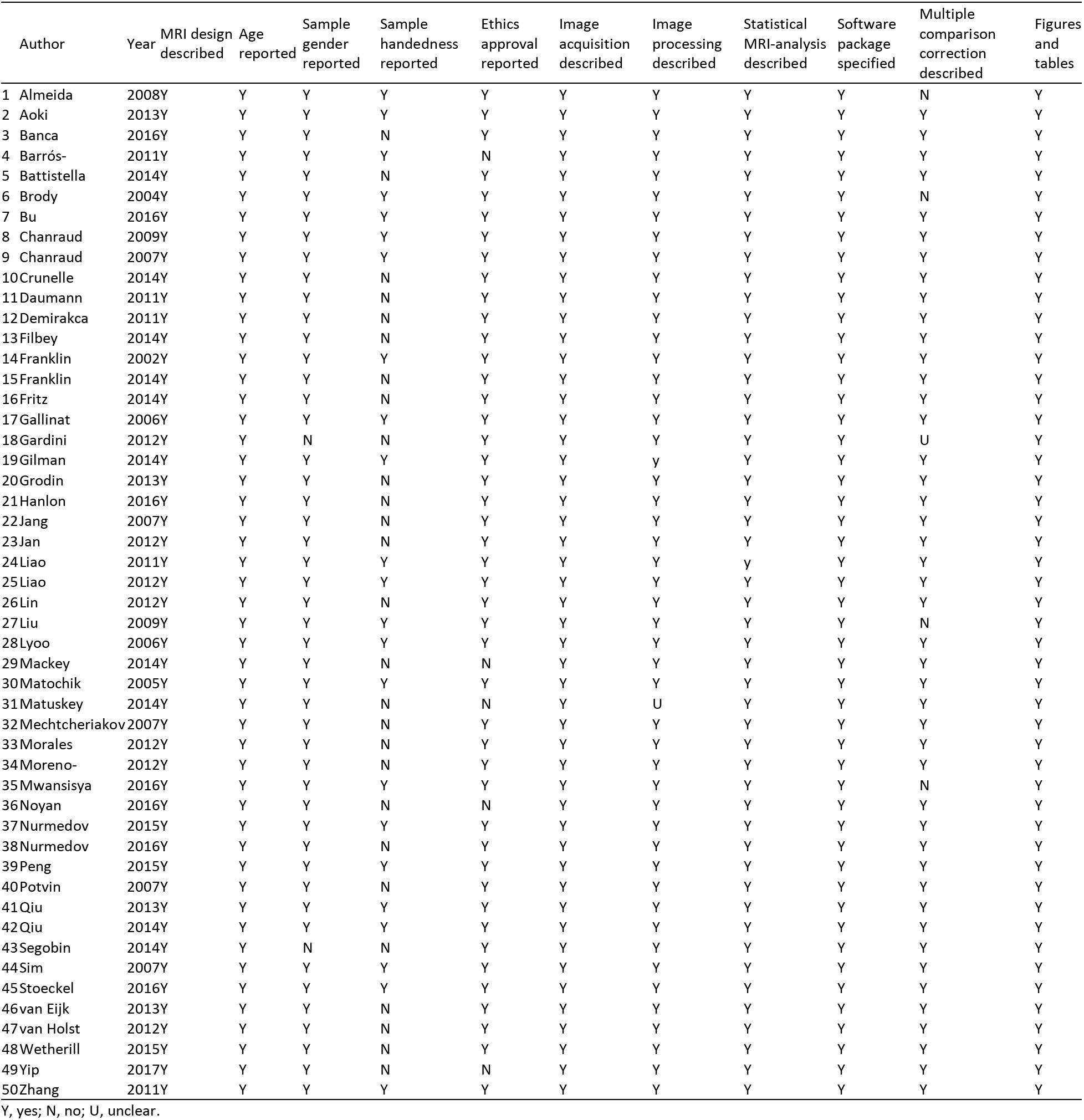
Summary of MRI quality.

| Author | Year | MRI design described | Age reported | Sample gender reported | Sample handedness reported | Ethics approval reported | Image acquisition described | Image processing described | Statistical MRI-analysis described | Software package specified | Multiple comparison correction described | Figures and tables |
| --- | --- | --- | --- | --- | --- | --- | --- | --- | --- | --- | --- | --- |
| 1 Almeida | 2008Y | Y | Y | Y | Y | Y | Y | Y | Y | Y | N | Y |
| 2 Aoki | 2013Y | Y | Y | Y | Y | Y | Y | Y | Y | Y | Y | Y |
| 3 Banca | 2016Y | Y | Y | N | Y | Y | Y | Y | Y | Y | Y | Y |
| 4 Barrós- | 2011Y | Y | Y | Y | Y | N | Y | Y | Y | Y | Y | Y |
| 5 Battistella | 2014Y | Y | Y | N | Y | Y | Y | Y | Y | Y | Y | Y |
| 6 Brody | 2004Y | Y | Y | Y | Y | Y | Y | Y | Y | Y | N | Y |
| 7 Bu | 2016Y | Y | Y | Y | Y | Y | Y | Y | Y | Y | Y | Y |
| 8 Chanraud | 2009Y | Y | Y | Y | Y | Y | Y | Y | Y | Y | Y | Y |
| 9 Chanraud | 2007Y | Y | Y | Y | Y | Y | Y | Y | Y | Y | Y | Y |
| 10Crunelle | 2014Y | Y | Y | N | Y | Y | Y | Y | Y | Y | Y | Y |
| 11Daumann | 2011Y | Y | Y | N | Y | Y | Y | Y | Y | Y | Y | Y |
| 12Demirakca | 2011Y | Y | Y | N | Y | Y | Y | Y | Y | Y | Y | Y |
| 13Filbey | 2014Y | Y | Y | N | Y | Y | Y | Y | Y | Y | Y | Y |
| 14Franklin | 2002Y | Y | Y | Y | Y | Y | Y | Y | Y | Y | Y | Y |
| 15Franklin | 2014Y | Y | Y | N | Y | Y | Y | Y | Y | Y | Y | Y |
| 16Fritz | 2014Y | Y | Y | N | Y | Y | Y | Y | Y | Y | Y | Y |
| 17Gallinat | 2006Y | Y | Y | Y | Y | Y | Y | Y | Y | Y | Y | Y |
| 18Gardini | 2012Y | Y | N | N | Y | Y | Y | Y | Y | Y | U | Y |
| 19Gilman | 2014Y | Y | Y | Y | Y | Y | Y | Y | Y | Y | Y | Y |
| 20Grodin | 2013Y | Y | Y | N | Y | Y | Y | Y | Y | Y | Y | Y |
| 21Hanlon | 2016Y | Y | Y | N | Y | Y | Y | Y | Y | Y | Y | Y |
| 22Jang | 2007Y | Y | Y | N | Y | Y | Y | Y | Y | Y | Y | Y |
| 23Jan | 2012Y | Y | Y | N | Y | Y | Y | Y | Y | Y | Y | Y |
| 24Liao | 2011Y | Y | Y | Y | Y | Y | Y | Y | Y | Y | Y | Y |
| 25Liao | 2012Y | Y | Y | Y | Y | Y | Y | Y | Y | Y | Y | Y |
| 26Lin | 2012Y | Y | Y | N | Y | Y | Y | Y | Y | Y | Y | Y |
| 27Liu | 2009Y | Y | Y | Y | Y | Y | Y | Y | Y | Y | N | Y |
| 28Lyoo | 2006Y | Y | Y | Y | Y | Y | Y | Y | Y | Y | Y | Y |
| 29Mackey | 2014Y | Y | Y | N | N | Y | Y | Y | Y | Y | Y | Y |
| 30Matochik | 2005Y | Y | Y | Y | Y | Y | Y | Y | Y | Y | Y | Y |
| 31Matuskey | 2014Y | Y | Y | N | N | Y | U | Y | Y | Y | Y | Y |
| 32Mechtcheriakov | 2007Y | Y | Y | N | Y | Y | Y | Y | Y | Y | Y | Y |
| 33Morales | 2012Y | Y | Y | N | Y | Y | Y | Y | Y | Y | Y | Y |
| 34Moreno- | 2012Y | Y | Y | N | Y | Y | Y | Y | Y | Y | Y | Y |
| 35Mwansisya | 2016Y | Y | Y | Y | Y | Y | Y | Y | Y | Y | N | Y |
| 36Noyan | 2016Y | Y | Y | N | N | Y | Y | Y | Y | Y | Y | Y |
| 37Nurmedov | 2015Y | Y | Y | Y | Y | Y | Y | Y | Y | Y | Y | Y |
| 38Nurmedov | 2016Y | Y | Y | N | Y | Y | Y | Y | Y | Y | Y | Y |
| 39Peng | 2015Y | Y | Y | Y | Y | Y | Y | Y | Y | Y | Y | Y |
| 40Potvin | 2007Y | Y | Y | N | Y | Y | Y | Y | Y | Y | Y | Y |
| 41Qiu | 2013Y | Y | Y | Y | Y | Y | Y | Y | Y | Y | Y | Y |
| 42Qiu | 2014Y | Y | Y | Y | Y | Y | Y | Y | Y | Y | Y | Y |
| 43Segobin | 2014Y | Y | N | N | Y | Y | Y | Y | Y | Y | Y | Y |
| 44Sim | 2007Y | Y | Y | Y | Y | Y | Y | Y | Y | Y | Y | Y |
| 45Stoeckel | 2016Y | Y | Y | Y | Y | Y | Y | Y | Y | Y | Y | Y |
| 46van Eijk | 2013Y | Y | Y | N | Y | Y | Y | Y | Y | Y | Y | Y |
| 47van Holst | 2012Y | Y | Y | N | Y | Y | Y | Y | Y | Y | Y | Y |
| 48Wetherill | 2015Y | Y | Y | N | Y | Y | Y | Y | Y | Y | Y | Y |
| 49Yip | 2017Y | Y | Y | N | N | Y | Y | Y | Y | Y | Y | Y |
| 50Zhang | 2011Y | Y | Y | Y | Y | Y | Y | Y | Y | Y | Y | Y |

Y, yes; N, no; U, unclear.

**Supplementary Table 3.**
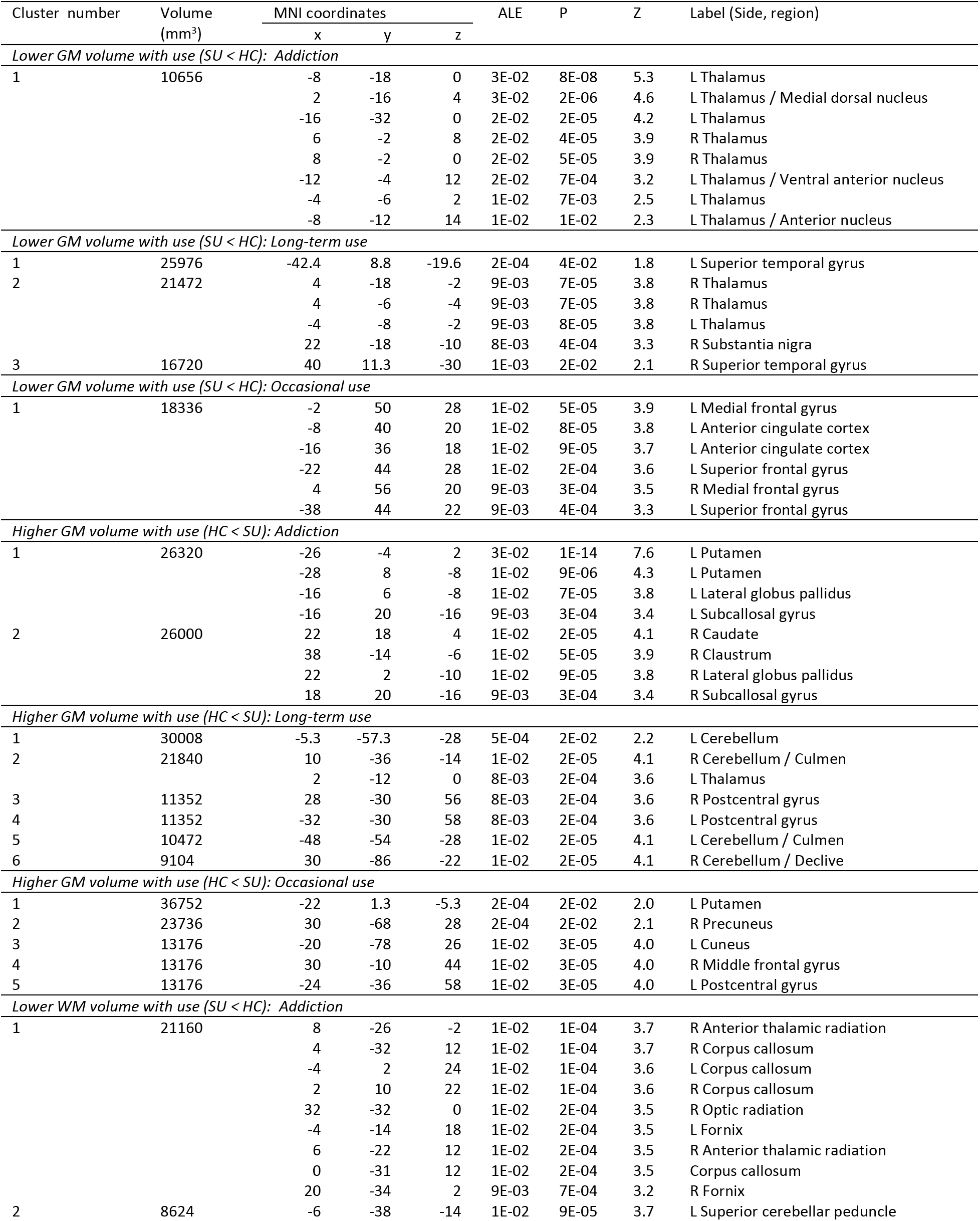
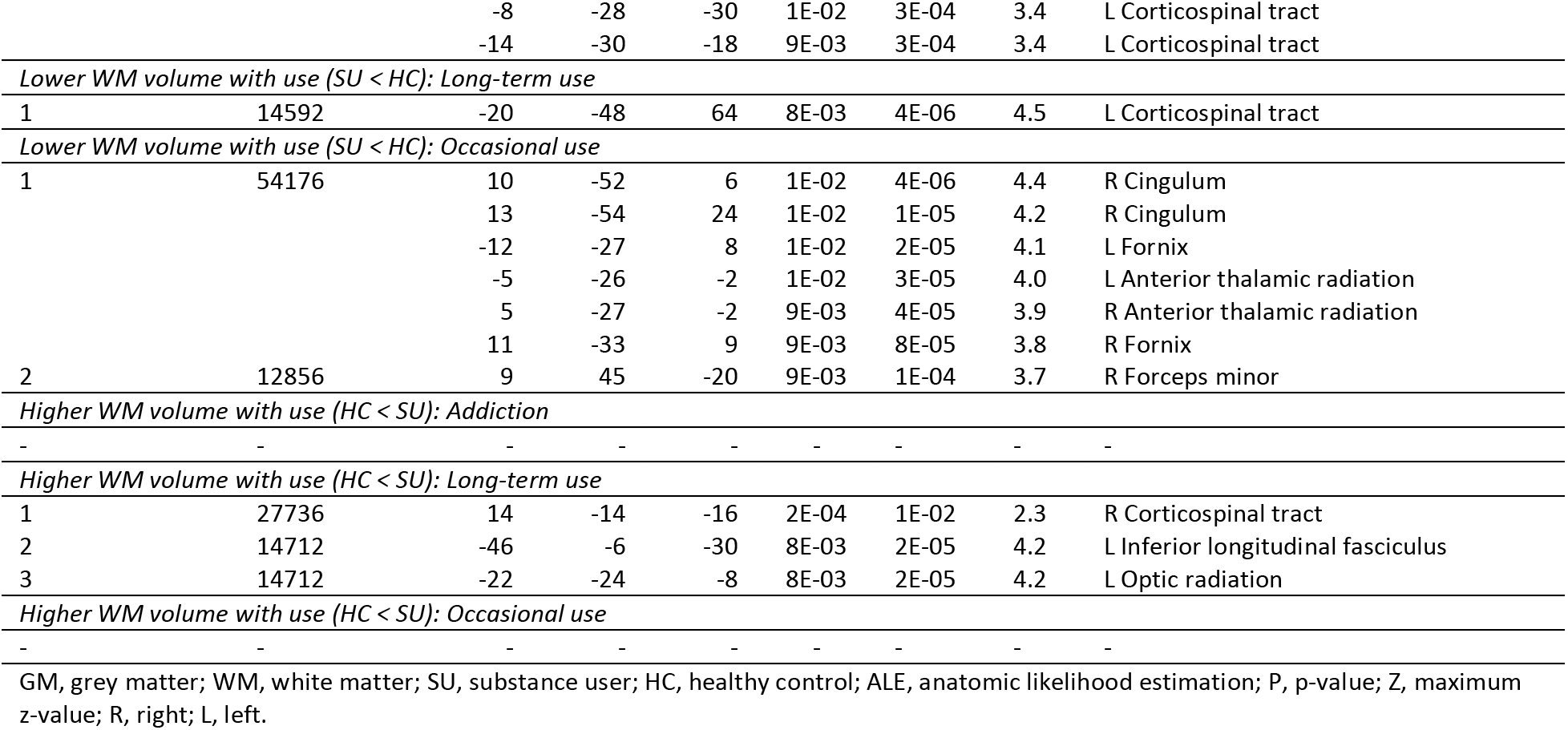
Type of consumption subgroup anatomic likelihood estimation meta-analytic results for studies comparing brain morphological changes between SU and HC (all substances), at cluster level inference p << 0.05 (FWE).

**Supplementary Table 4.**
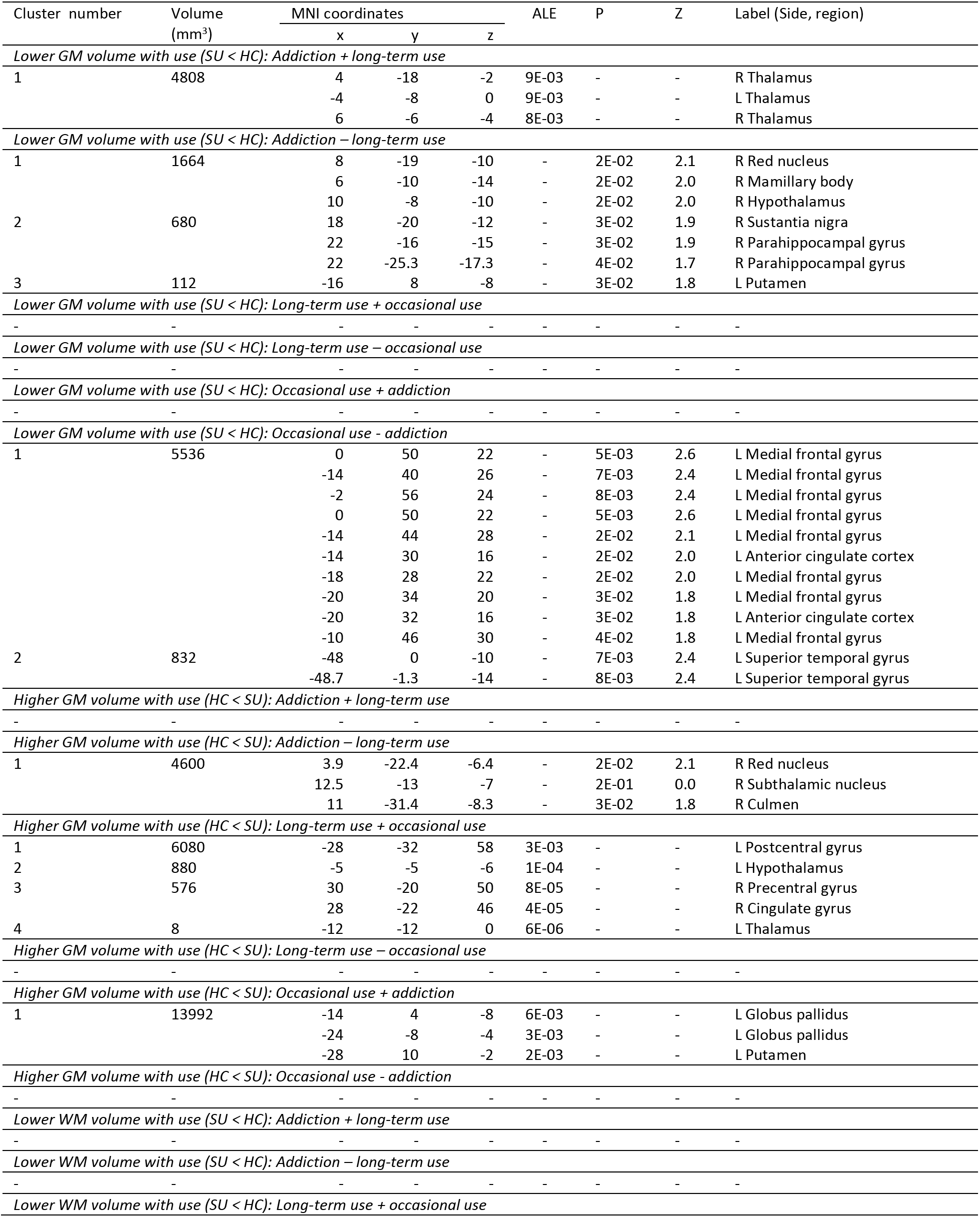
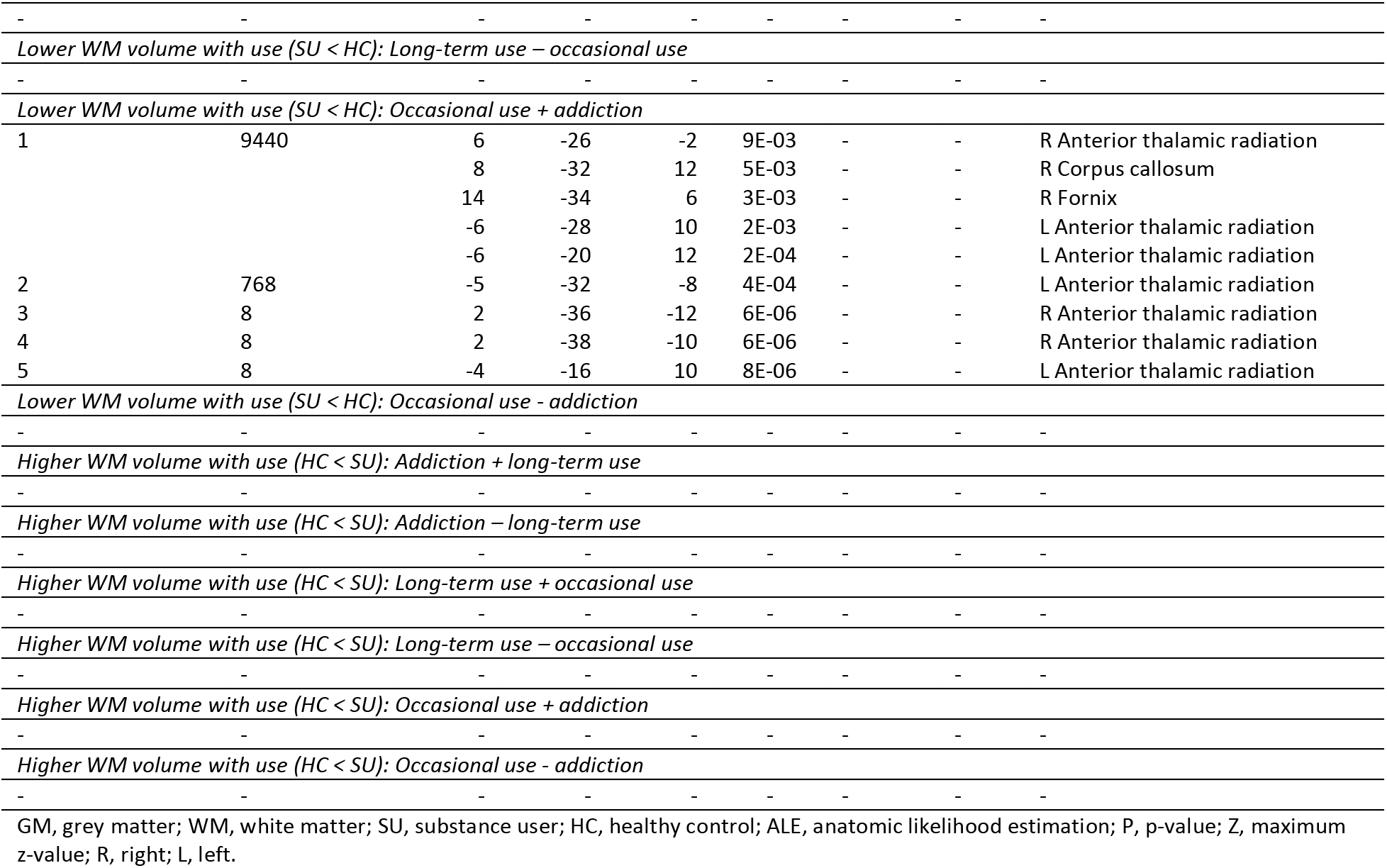
Contrast analysis of type of consumption subgroup anatomic likelihood estimation meta-analytic results for studies comparing brain morphological changes between SU and HC (all substances), at cluster level inference p < 0.05 (FWE).

**Supplementary Table 5.**
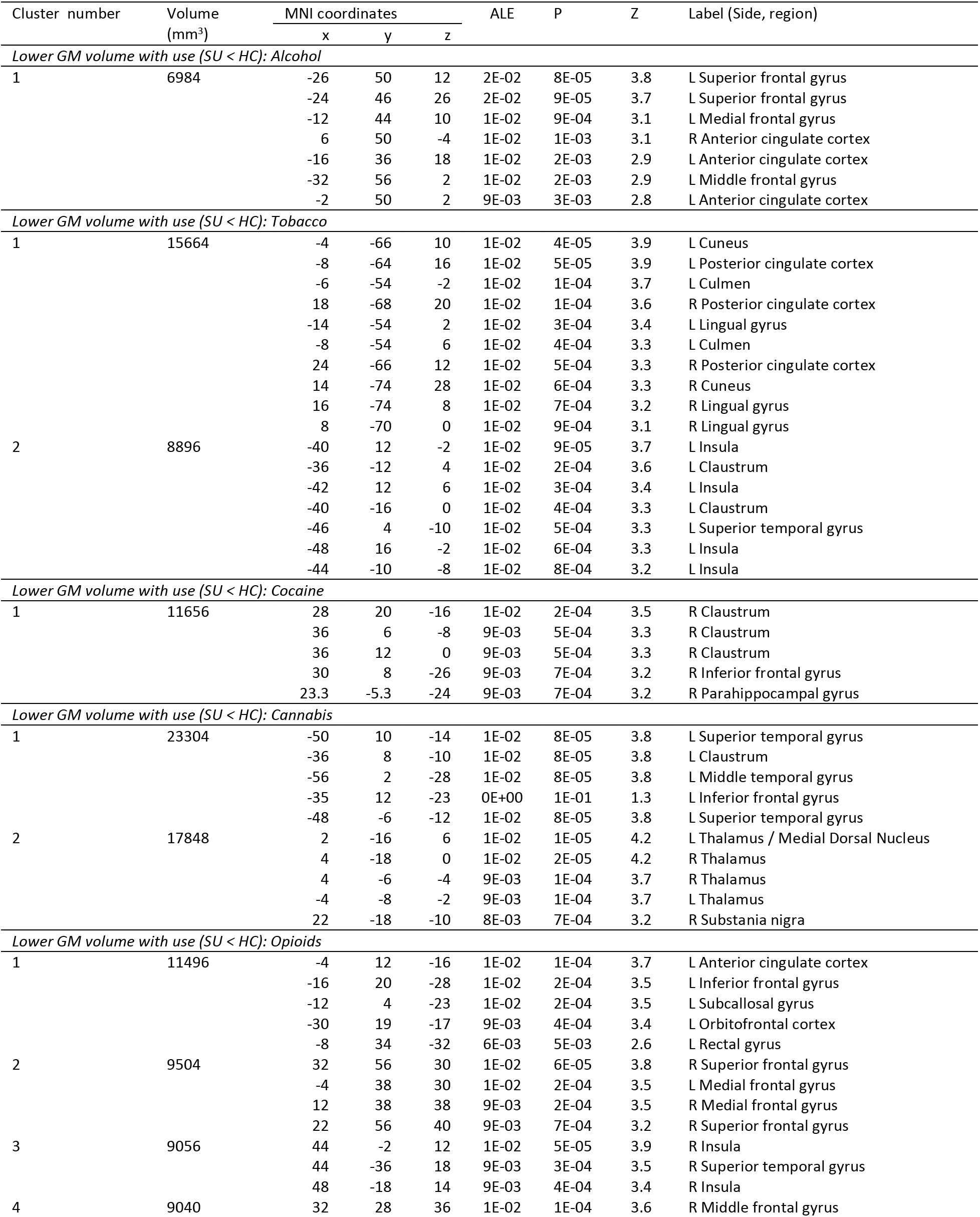
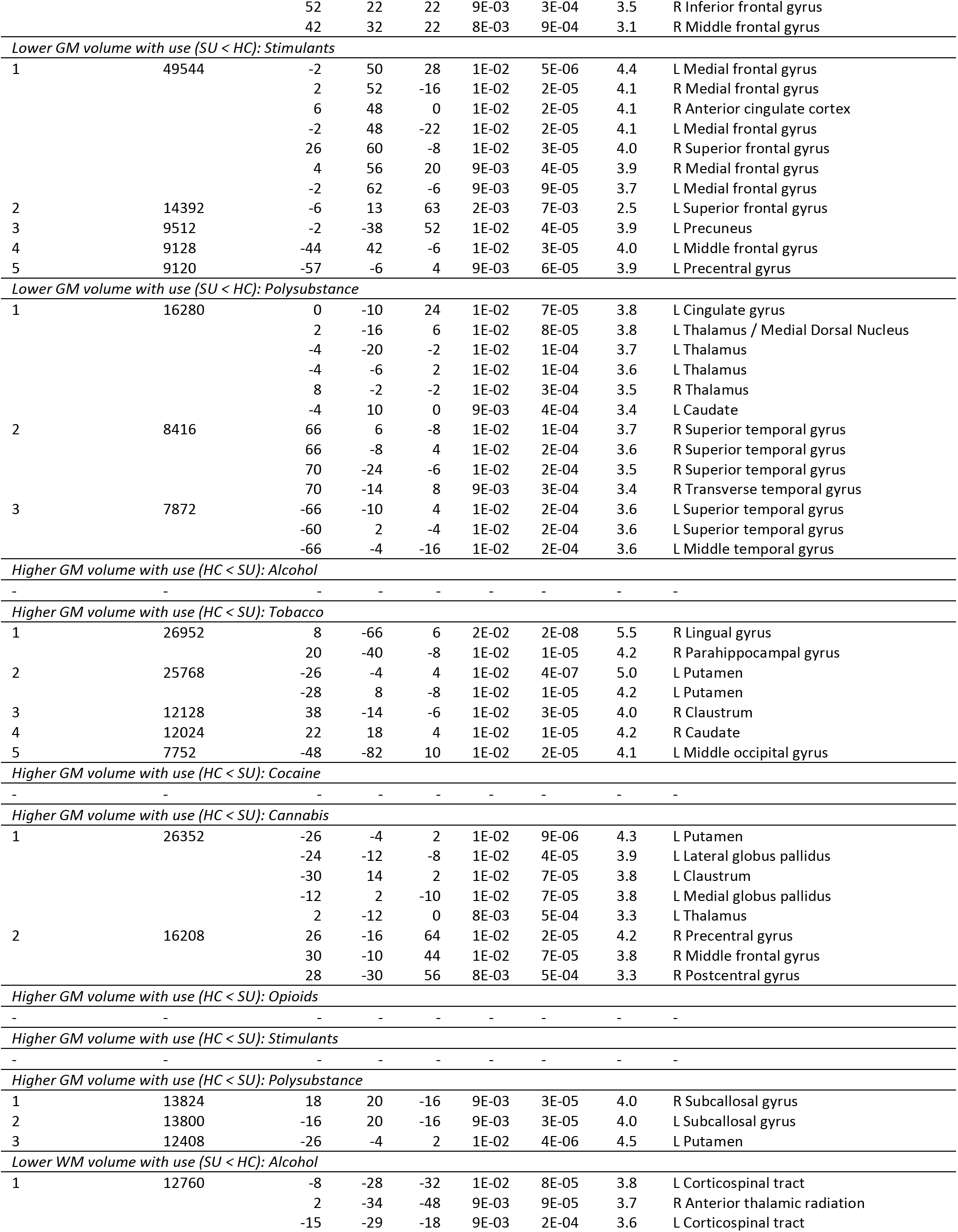
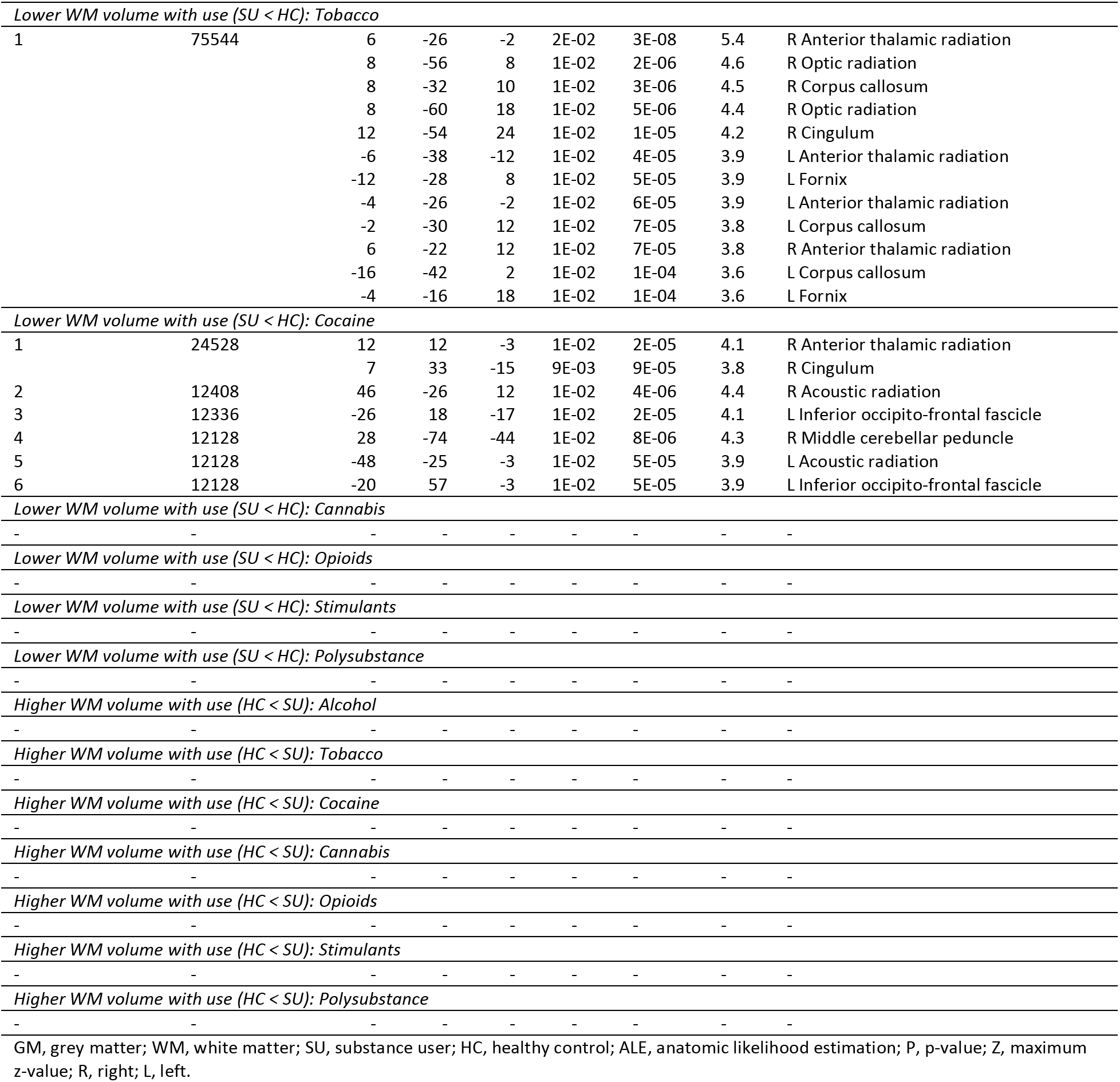
Type of substance subgroup anatomic likelihood estimation meta-analytic results for studies comparing brain morphological changes between SU and HC, at cluster level inference p < 0.05 (FWE).

**Supplementary Table 6.**
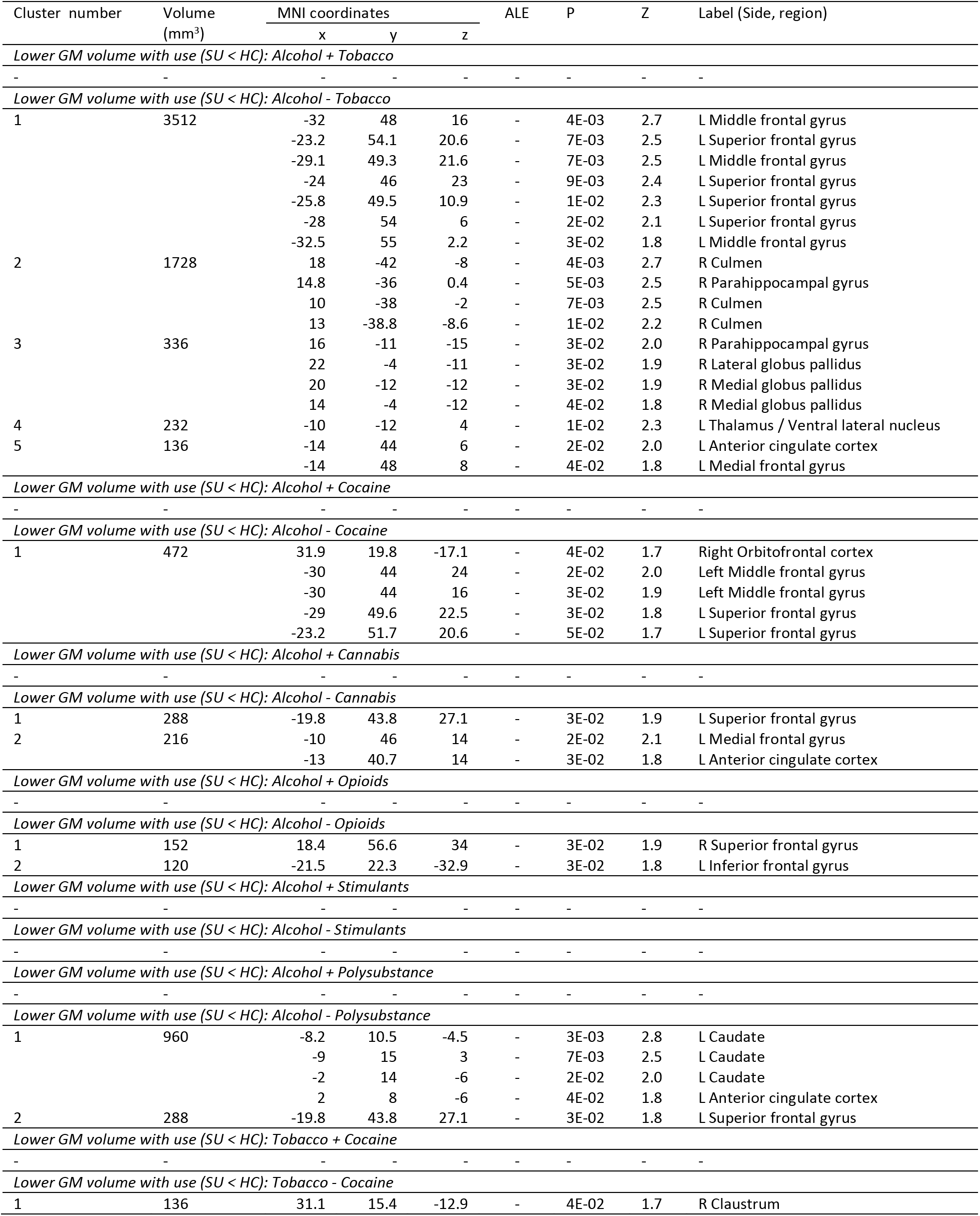
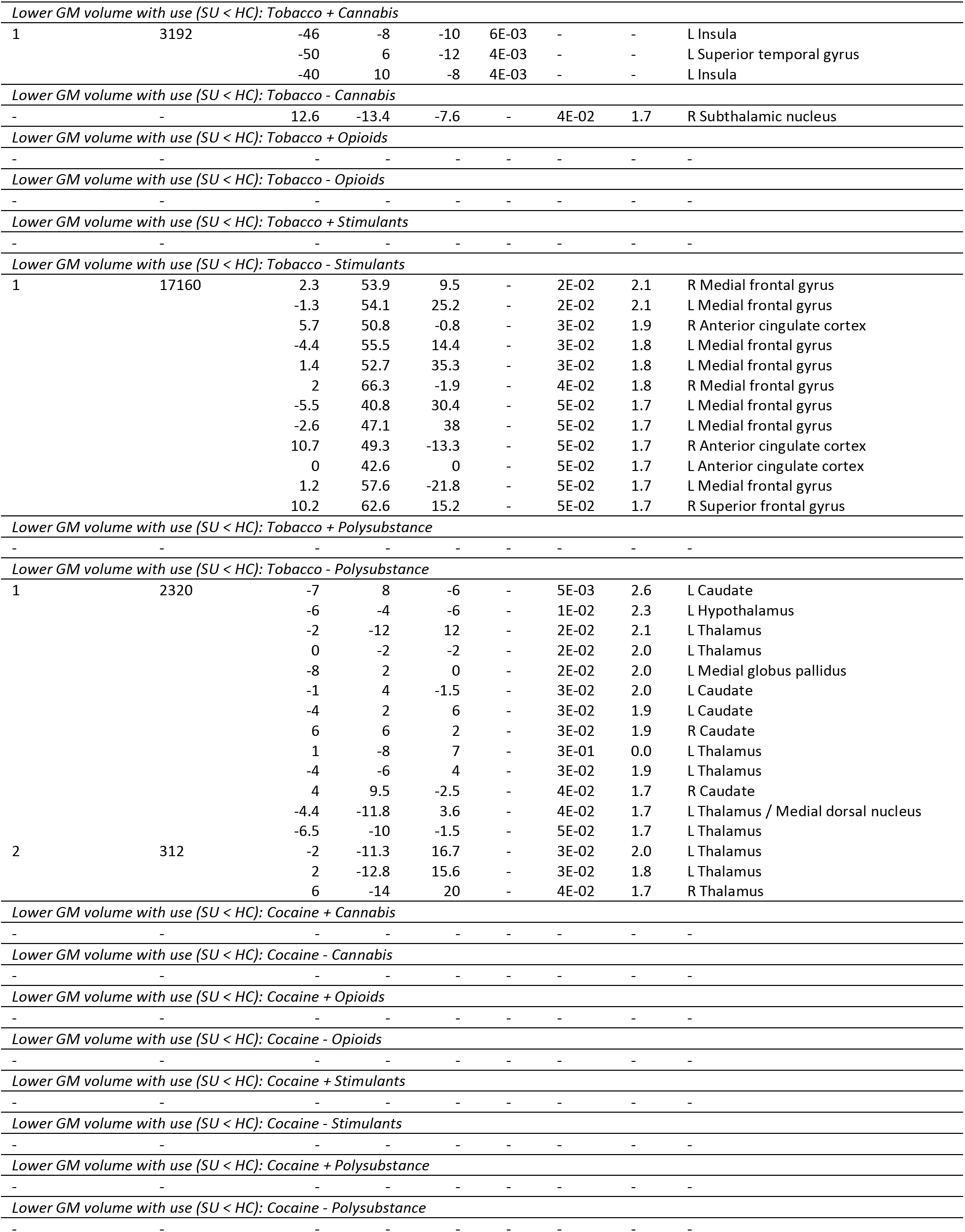
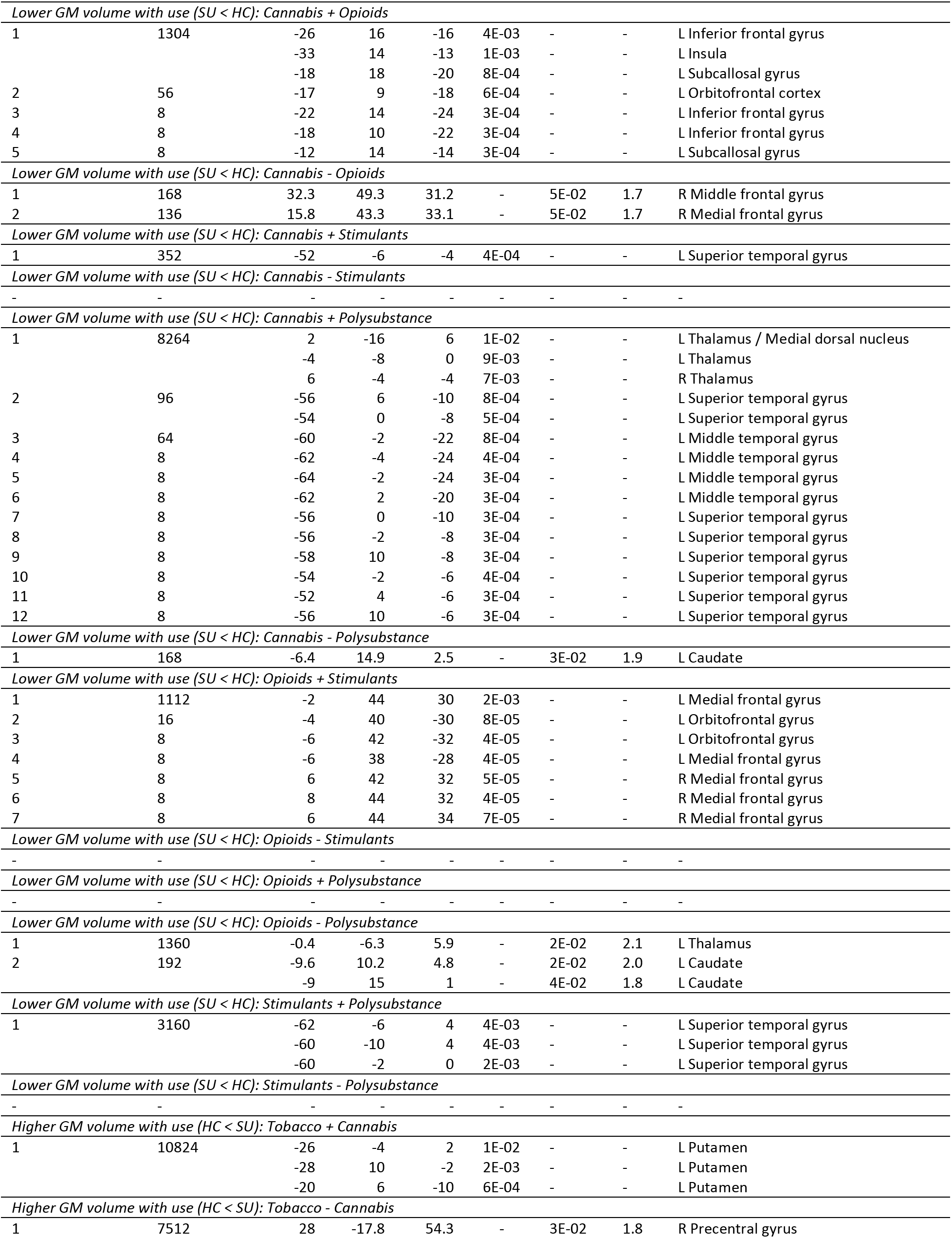
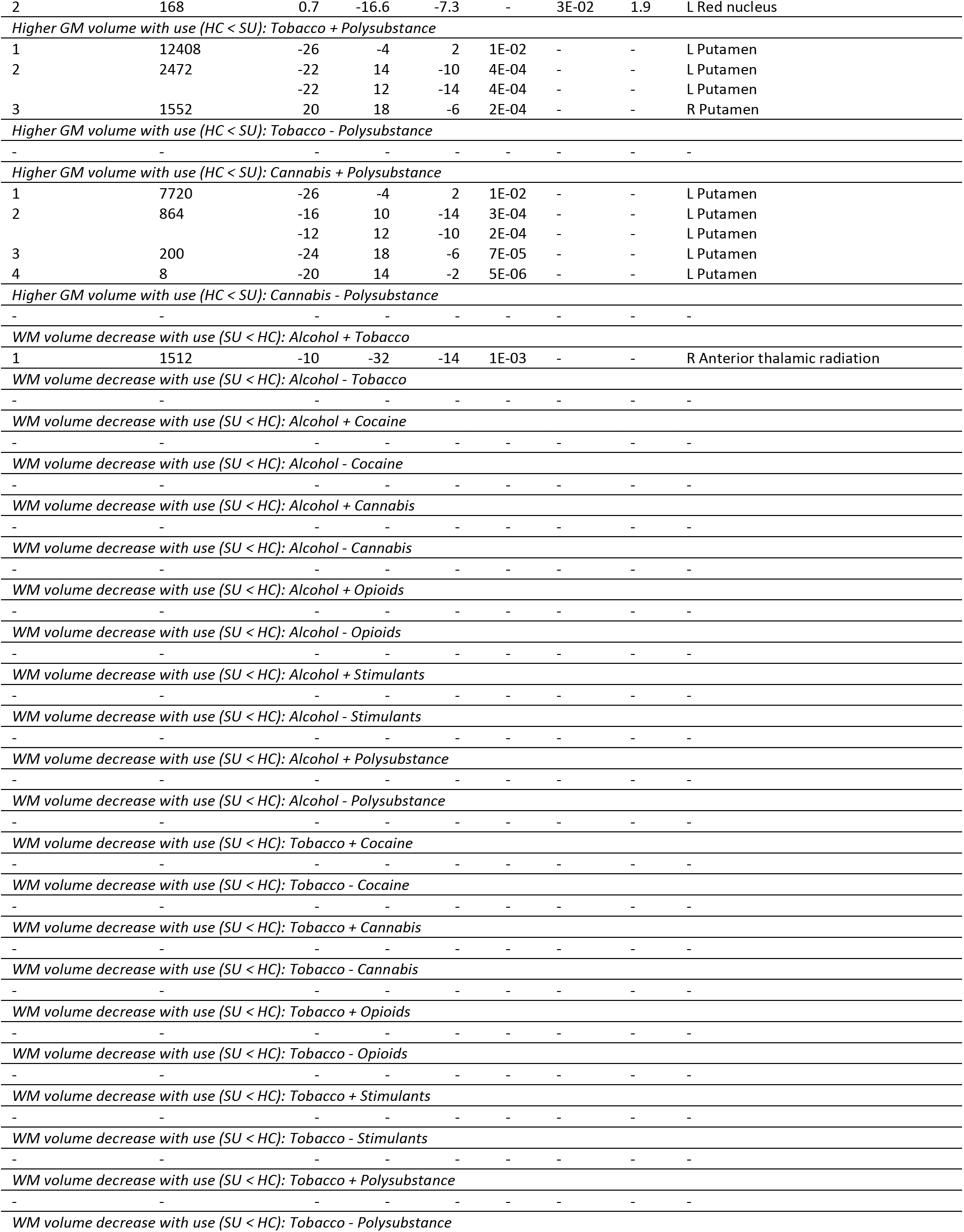
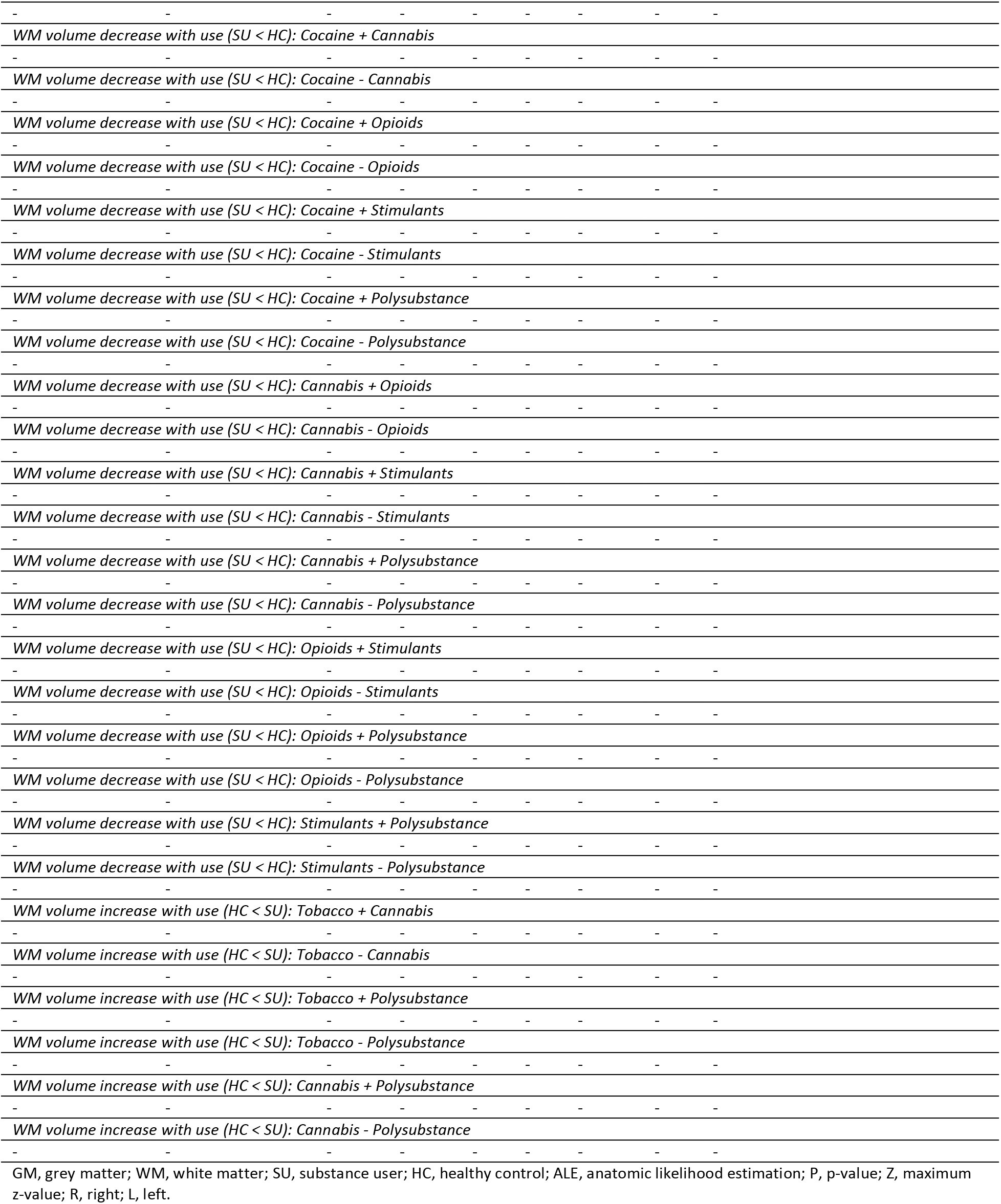
Contrast analysis of type of substance subgroup anatomic likelihood estimation meta-analytic results for studies comparing brain morphological changes between SU and HC, at cluster level inference p < 0.05 (FWE).

**Supplementary Table 7.**
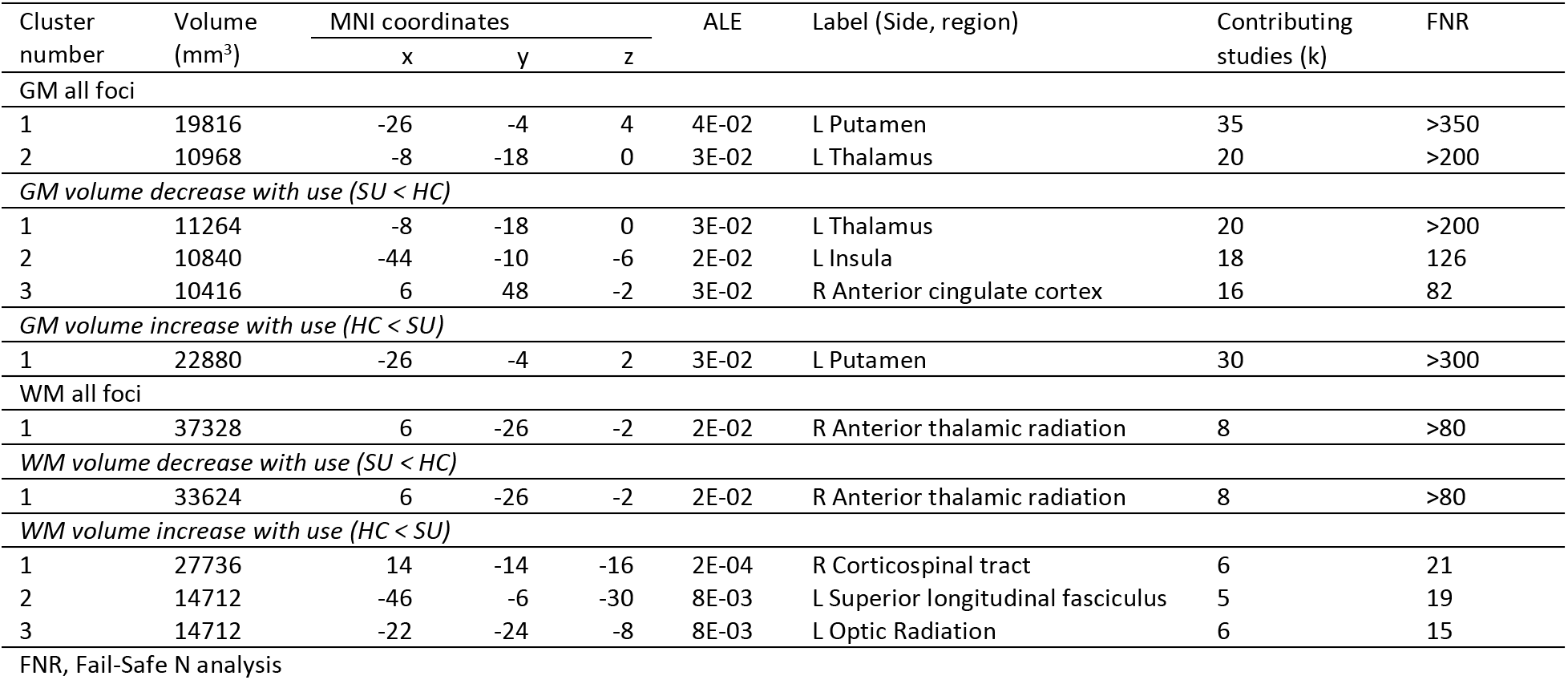
FNR robustness assessment for significant ALE maps resulting from primary outcomes.

